# UNC-6/Netrin and its Receptors UNC-5 and UNC-40/DCC Control Growth Cone Polarity, Microtubule Accumulation, and Protrusion

**DOI:** 10.1101/294215

**Authors:** Mahekta R. Gujar, Lakshmi Sundararajan, Aubrie Stricker, Erik A. Lundquist

## Abstract

Many axon guidance ligands and their receptors have been identified, but it is still unclear how these ligand-receptor interactions regulate events in the growth cone, such as protrusion and cytoskeletal arrangement, during directed outgrowth *in vivo*. In this work, we dissect the multiple and complex effects of UNC-6/Netrin on the growth cone. Previous studies showed that in *C. elegans*, the UNC-6/Netrin receptor UNC-5 regulates growth cone polarity, as evidenced by loss of asymmetric dorsal F-actin localization and protrusion in *unc-5* mutants. UNC-5 and another UNC-6/Netrin receptor UNC-40/DCC also regulate the extent of protrusion, with UNC-40/DCC driving protrusion and UNC-5 inhibiting protrusion. In this work we analyze the roles of UNC-6/Netrin, UNC-40/DCC, and UNC-5 in coordinating growth cone F-actin localization, microtubule organization, and protrusion that results in directed outgrowth away from UNC-6/Netrin. We find that a previously-described pathway involving the UNC-73/Trio Rac GEF and UNC-33/CRMP that acts downstream of UNC-5, regulates growth cone dorsal asymmetric F-actin accumulation and protrusion. *unc-5* and *unc-33* mutants also display excess EBP-2::GFP puncta, suggesting that MT + end accumulation is important in growth cone polarity and/or protrusion. *unc-73* Rac GEF mutants did not display excess EBP-2::GFP puncta despite larger and more protrusive growth cones, indicating a MT-independent mechanism to polarize the growth cone and to inhibit protrusion, possibly via actin. Finally, we show that UNC-6/Netrin and UNC-40/DCC are required for excess protrusion in *unc-5* mutants, but not for loss of F-actin asymmetry or MT + end accumulation, indicating that UNC-6/Netrin and UNC-40/DCC are required for protrusion downstream of F-actin asymmetry and MT + end entry. Our data suggest a model in which UNC-6/Netrin polarizes the growth cone via UNC-5, and then regulates a balance of pro- and anti-protrusive forces driven by UNC-40 and UNC-5, respectively, that result in directed protrusion and outgrowth.

## Introduction

Neural circuits and networks are formed by intricate interactions of axonal growth cones with the extracellular environment (Tessier-Lavigne and Goodman 1996; Mortimer *et al.* 2008). Many extracellular molecules that guide growth cone migrations have been identified, but the effects of these guidance molecules on growth cone morphology during outgrowth *in vivo* are incompletely understood.

The secreted UNC-6/Netrin guidance cue and its receptors UNC-5 and UNC-40/DCC guide cell and growth cone migrations in a manner conserved from invertebrates to mammals. In *C. elegans*, UNC-6/Netrin is expressed in cells along the ventral midline, including neurons with axons that extend down the lengths of the ventral nerve cord (Wadsworth *et al.* 1996; Asakura *et al.* 2007). UNC-6 controls both ventral migrations (towards UNC-6) and dorsal migrations (away from UNC-6), and *unc-6* mutants have defects in both ventral and dorsal guidance (Hedgecock *et al.* 1990; Norris and Lundquist 2011). Ventral versus dorsal responses to UNC-6/Netrin are mediated by expression of UNC-40 and UNC-5 on growth cones. Classically, UNC-40 was thought to mediate ventral growth toward UNC-6 (Chan *et al.* 1996), and UNC-5 was thought to mediate dorsal growth away from UNC-6 (Leung-Hagesteijn 1992), although UNC-40 also acts in dorsal growth along with UNC-5, likely as a heterodimer (Hong *et al.* 1999; Macneil *et al.* 2009; Norris and Lundquist 2011; Norris *et al.* 2014). Recent studies indicate that UNC-5 can act also act in ventral migrations (Levy-Strumpf and Culotti 2014; Yang *et al.* 2014; Limerick *et al.* 2018), and might serve to focus UNC-40 localization ventrally in the cell body toward the UNC-6/Netrin source. Thus, the roles of UNC-40 and UNC-5 in ventral and dorsal growth are more complex than initially appreciated.

The growth cones of the VD motor neuron processes migrate dorsally in a commissural route to the dorsal nerve cord (Knobel *et al.* 1999; Norris and Lundquist 2011). The VD cell bodies reside in the ventral nerve cord, and extend processes anteriorly, which then turn and begin dorsal commissural migration away from UNC-6. Mutations in *unc-6, unc-5,* and *unc-40* disrupt the dorsal guidance of the VD axons (Hedgecock *et al.* 1990). Commissural VD growth cones display robust and dynamic lamellipodial and filopodial protrusion localized to the dorsal leading edge, away from the UNC-6/Netrin source, resulting in directed dorsal migration (Knobel *et al.* 1999; Norris and Lundquist 2011). F-actin also accumulated at the dorsal leading edge, near the site of protrusion (Norris and Lundquist 2011). Previous studies showed that UNC-6/Netrin, UNC-5, and UNC-40/DCC control lamellipodial and filopodial protrusion of VD growth cones (Norris and Lundquist 2011; Norris *et al.* 2014). *unc-5* mutant VD growth cones showed excess protrusion, with larger lamellipodial growth cone bodies and longer and longer-lasting filopodial protrusions. Furthermore, the protrusions were no longer focused to the dorsal leading edge but occurred all around the growth cone. Finally, F-actin was no longer restricted to the dorsal leading edge but was found throughout the periphery of the growth cone in *unc-5* mutants. Thus, these large, unfocused *unc-5* mutant growth cones moved very little, consistent with findings in cultured growth cones that large, more protrusive growth cones exhibited reduced rates of movement (Ren and Suter 2016).

While the effect of *unc-5* mutation on VD growth cones was severe, loss of *unc-40* had no significant effect on extent or polarity of protrusion (Norris and Lundquist 2011). However, constitutive activation of UNC-40 signaling (MYR::UNC-40) in VD growth cones led to small growth cones with little or no protrusion, similar to constitutive activation of UNC-5 (MYR::UNC-5) (Norris and Lundquist 2011; Norris *et al.* 2014). Functional UNC-5 was required for the inhibitory effects of MYR::UNC-40. This suggests that MYR::UNC-40 acts as a heterodimer with UNC-5 to inhibit protrusion, and in *unc-40* mutants, UNC-5 alone was sufficient to inhibit protrusion. *unc-6(ev400)* null mutants had no effect on extent of VD growth cone protrusion, but did affect polarity of protrusion as well as F-actin polarity, both lost in *unc-6* mutants (Norris and Lundquist 2011). Thus, UNC-6/Netrin affects VD growth cone polarity (F-actin and protrusion), but it is unclear if the effects on extent of protrusion by UNC-5 and UNC-40 involve UNC-6/Netrin.

These results suggest that in the same VD growth cone, UNC-40 can both drive protrusion and inhibit protrusion along with UNC-5. That the normal VD growth cone has polarized protrusion to the dorsal leading edge and reduced protrusion ventrally near the axon shaft suggests that the activities of UNC-40 and UNC-40-UNC-5 might be asymmetric across the growth cone. The *unc-6(e78)* mutation, which specifically affects interaction with UNC-5, causes excess growth cone protrusion and abolished polarity similar to *unc-5* mutants (Norris and Lundquist 2011), suggesting that UNC-6/Netrin inhibits protrusion through UNC-5. The involvement of UNC-6/Netrin in pro-protrusive UNC-40 activity is unclear.

UNC-5 affects three aspects of growth cone morphology during outgrowth *in vivo*: polarity of protrusion to the dorsal leading edge, F-actin asymmetric accumulation to the dorsal leading edge, and inhibition of growth cone protrusion (Norris and Lundquist 2011; Norris *et al.* 2014). Growth cone motility and guidance is dependent on the actin and microtubule cytoskeleton (Dent and Gertler 2003). The axon shaft and the central region of the growth cone is composed of bundled microtubules with their plus (+) ends (MT+) oriented towards the growth cone. The peripheral region of the growth cone contains highly dynamic actin that is relatively free of microtubules. The actin filaments at the leading edge of the growth cone form a branched lamellipodial meshwork and filopodial bundles in the essential in sensing guidance cues and driving the forward motion of the axon (Forscher and Smith 1988; Gallo and Letourneau 2004; Pak *et al.* 2008; Dent *et al.* 2011; Omotade *et al.* 2017).

Our previous results show an effect of UNC-5 on F-actin polarity and protrusive events driven by F-actin. We recently showed that a family of genes encoding flavin monooxygenases (FMOs) are required for inhibition of protrusion mediated by UNC-5 (Gujar *et al.* 2017). In Drosophila and mammals, the FMO-containing MICAL molecule causes actin depolymerization and collapse through direct oxidation of actin (Terman *et al.* 2002; Hung *et al.* 2010; Hung *et al.* 2011). We previously described a second signaling pathway downstream of UNC-5 required to inhibit protrusion that includes the UNC-73/Trio Rac GEF, the Rac GTPases CED-10 and MIG-2, and the UNC-33/CRMP cytoskeletal molecule (Norris *et al.* 2014), which can interact with MTs in other systems (Fukata *et al.* 2002). In cultured growth cones, most stable microtubules remain in the central domain. A small population of dynamic microtubules can explore the periphery and penetrate the filopodia, where they interact with extracellular cues resulting in proper axonal elongation and guidance (Sabry *et al.* 1991; Tanaka *et al.* 1995; Dent and Gertler 2003; Lowery and Van Vactor 2009). Thus, motility of the growth cone is achieved through proper regulation and coordination between microtubules and the actin cytoskeleton (Dent and Kalil 2001; Buck and Zheng 2002; Zhou *et al.* 2002; Zhou and Cohan 2004). MTs have been implicated in Unc5 signaling (Shao *et al.* 2017; Huang *et al.* 2018), but the role of MTs in UNC-5-mediated VD growth cone outgrowth *in vivo* remains unclear.

That UNC-5 and UNC-40 cooperate to guide migrations of axons that grow toward UNC-6/Netrin indicates that the roles of these molecules are more complex than discrete “attractive” and “repulsive” functions. The signaling pathways used by UNC-40 are well-described, but the intracellular pathways used by UNC-5 remain unclear. In this work, we analyze three aspects of growth cone behavior to understand the roles of these molecules in growth away from UNC-6/Netrin: growth cone protrusion; F-actin asymmetric accumulation; and EBP-2::GFP distribution, which has been used previously to monitor MT + ends in *C. elegans* embryos and neurons (Srayko *et al.* 2005; Kozlowski *et al.* 2007; Yan *et al.* 2013). We find that UNC-6/Netrin is required for the excess protrusion in *unc-5* mutant growth cones, similar to UNC-40, and that UNC-6/Netrin, along with UNC-5, polarizes growth cone F-actin accumulation and protrusion to the dorsal leading edge, resulting in focused dorsal protrusion of the growth cone. We find that EBP-2::GFP puncta are found in excess in *unc-5, unc-6,* and *unc-33* mutant growth cones, suggesting that UNC-6/Netrin and UNC-5 signaling can block MT + end accumulation in growth cones, which correlates with inhibited growth cone protrusion, and suggests a pro-protrusive role for MTs in the growth cone. Finally, we show that UNC-6/Netrin and UNC-40 stimulate VD growth cone protrusion downstream of dorsal F-actin polarity and growth cone EBP-2::GFP accumulation. An implication of our results is that in a growth cone growing away from an UNC-6/Netrin source, UNC-6/Netrin both stimulates protrusion dorsally, away from the source, and inhibits protrusion ventrally, near the source, resulting in directed outgrowth.

## Materials and Methods

### Genetic methods

Experiments were performed at 20°C using standard *C. elegans* techniques. Mutations used were LGI: *unc-40(n324* and *e1430)*, *unc-73(rh40, e936, ev802* and *ce362)*; LGII: *juIs76[Punc-25::gfp]*. LGIV: *unc-5(e53, e553, e791* and *e152)*, *unc-33(e204* and *e1193*), *unc-44(e362, e1197* and *e1260*), *ced-10(n1993)*; LGX: *unc-6(ev400), unc-6(e78), mig-2(mu28), lqIs182* [*Punc-25::mig-2(G16V)*]*, lqIs170* [*rgef-1::vab-10ABD::gfp*]. Chromosomal locations not determined: *lqIs279* and *lqIs280* [*Punc-25::ebp-2::gfp*], *lqIs296* [*Punc-25::myr::unc-5*]*, lhIs6* [*Punc-25::mCherry*]. The presence of mutations in single and double mutant strains was confirmed by phenotype, PCR genotyping, and sequencing.

Extrachromosomal arrays were generated using standard gonadal injection (Mello and Fire 1995) and include: *lqEx999* and *lqEx1000* [*Punc-25::myr::unc-40; Pgcy-32::yfp*], *lqEx1017* and *lqEx1018* [*Punc-25::ced-10(G12V); Pgcy-32::yfp*]. Multiple (≥3) extrachromosomal transgenic lines of transgenes described here were analyzed with similar effect, and one was chosen for integration and further analysis. The *mig-2(mu28); ced-10(n1993M+)* strain was balanced with the nT1 balancer. The *Punc-25::ebp-2::gfp* plasmid was constructed using standard recombinant DNA techniques. The sequences of all plasmids and all oligonucleotides used in their construction are available upon request.

### Growth cone imaging

VD growth cones were imaged and quantified as previously described (Norris and Lundquist 2011). Briefly, animals at ~16 h post-hatching at 20°C were placed on a 2% agarose pad and paralyzed with 5mM sodium azide in M9 buffer, which was allowed to evaporate for 4 min before placing a coverslip over the sample. Some genotypes were slower to develop than others, so the 16 h time point was adjusted for each genotype. Growth cones were imaged with a Qimaging Rolera mGi camera on a Leica DM5500 microscope. Images were analyzed in ImageJ, and statistical analyses done with Graphpad Prism software. As described in (Norris and Lundquist 2011; Norris *et al.* 2014), growth cone area was determined by tracing the perimeter of the growth cone body, not including filopodia. Average filopodial length was determined using a line tool to trace the length of the filopodium. Unless otherwise indicated, ≥25 growth cones were analyzed for each genotype. These data were gathered in ImageJ and entered into Graphpad Prism for analysis. A two-sided *t*-test with unequal variance was used to determine significance of difference between genotypes.

### VAB-10ABD::GFP imaging

The F-actin binding domain of VAB-10/spectraplakin fused to GFP has been used to monitor F-actin in *C. elegans* (Bosher *et al.* 2003; Patel *et al.* 2008). We used it to image F-actin in the VD growth cones as previously described (Norris and Lundquist 2011). To control for variability in growth cone size and shape, and as a reference for asymmetric localization of VAB-10ABD::GFP, a soluble mCherry volume marker was included in the strain. Growth cones images were captured as described above. ImageJ was used image analysis to determine asymmetric VAB-10ABG::GFP localization. For each growth cone, five line scans were made from dorsal to ventral (see Results). For each line, pixel intensity was plotted as a function of distance from the dorsal leading edge of the growth cone. The average intensity (arbitrary units) and standard error for each growth cone was determined. For dorsal versus ventral comparisons, the pixel intensities for VAB-10ABD::GFP were normalized to the volumetric mCherry fluorescence in line scans from the dorsal half and the ventral half of each growth cone. This normalized ratio was determined for multiple growth cones, and the average and standard error for multiple growth cones was determined. Statistical comparisons between genotypes were done using a two-tailed *t-*test with unequal variance on these average normalized ratios of multiple growth cones of each genotype.

### EBP-2::GFP imaging

EBP-2::GFP has previously been used to monitor microtubule plus ends in other *C. elegans* cells including neurons (Srayko *et al.* 2005; Kozlowski *et al.* 2007; Yan *et al.* 2013). We constructed a transgene consisting of the *unc-25* promoter driving expression of *ebp-2::gfp* in the VD/DD neurons. In growth cones, a faint fluorescence was observed throughout the growth cone, resembling a soluble GFP and allowing for the growth cone perimeter to be defined. In addition to this faint, uniform fluorescence, brighter puncta of EBP-2::GFP were observed that resembled the EBP-1::GFP puncta described in other cells and neurons. For each growth cone, the perimeter and filopodia were defined, and the EBP-2::GFP puncta in the growth cone were counted. For each genotype, the puncta number for many growth cones (≥25 unless otherwise noted) was determined. Puncta number displayed high variability within and between genotypes, so box-and-whiskers plots (Graphpad Prism) were used to accurately depict this variation. The grey boxes represent the upper and lower quartiles of the data set, and the “whiskers” represent the high and low values. Dots represent major outliers. Significance of difference was determined by a two-sided *t*-test with unequal variance.

### Data availability

Strains and plasmids are available upon request. The authors affirm that all data necessary for confirming the conclusions of the article are present within the article, figures, and tables.

## Results

### Functional UNC-6 is required for excess growth cone protrusion of *unc-5* mutants

Previous studies have shown that *unc-5* mutants displayed increased VD growth cone protrusiveness with larger growth cone area and longer filopodial protrusions as compared to wild-type (Norris and Lundquist 2011). However, *unc-5; unc-40* double mutants were found to have near wild-type levels of VD growth cone protrusion, suggesting that a functional UNC-40 was required for the over protrusive growth cone phenotype observed in *unc-5* loss of function mutants alone (Norris and Lundquist 2011). *unc-6(ev400)* also suppressed the excess growth cone protrusion (growth cone size and filopodial length) of *unc-5* mutants (Figure 1A-E), suggesting that excess protrusion in *unc-5* mutants also requires functional UNC-6.

**Figure 1.**
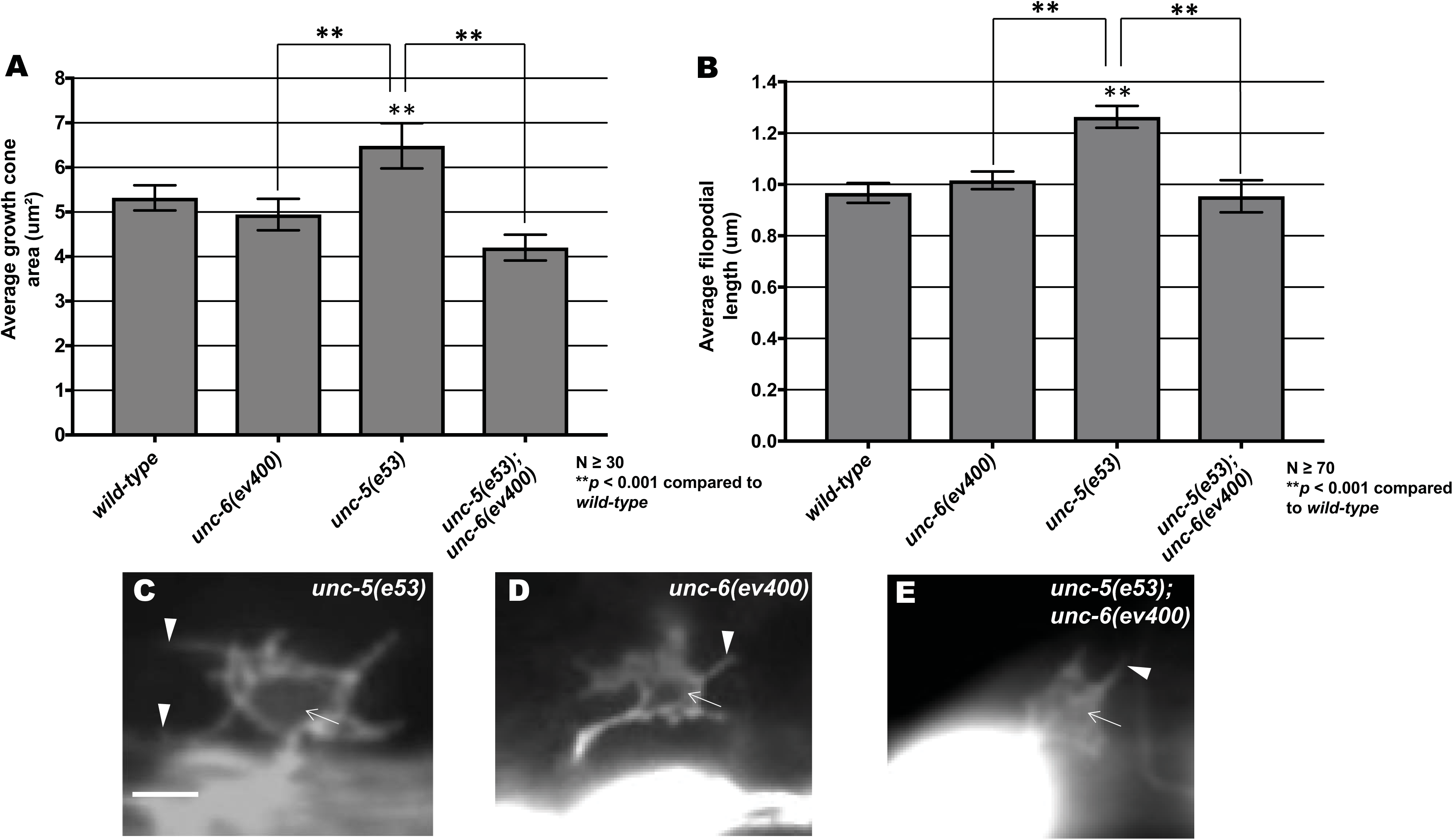
UNC-6 regulates growth cone protrusion. **(A-B)** Graphs of the average growth cone area and filopodial length in wild-type and mutants, as described in (Norris and Lundquist 2011) (See Methods). **(C-E)** Fluorescence micrographs of VD growth cones with *Punc-25::gfp* expression from the transgene *juIs76*. Arrows point to the growth cone body, and arrowheads to filopodial protrusions. Scale bar: 5μm.

### VD Growth cone F-actin and EBP-2::GFP organization

The F-actin binding domain of the spectraplakin VAB-10 was previously used to monitor F-actin in the VD growth cone in *C. elegans* (Norris and Lundquist 2011). In wild-type VD growth cones F-actin preferentially localized to the leading edge of the growth cone (~1.23 fold more accumulation in the dorsal half or the growth vs the ventral half) (Figure 2 and (Norris and Lundquist 2011)). Most growth cone filopodial protrusion occurs at the dorsal leading edge of the VD growth cone, correlating with F-actin accumulation.

**Figure 2.**
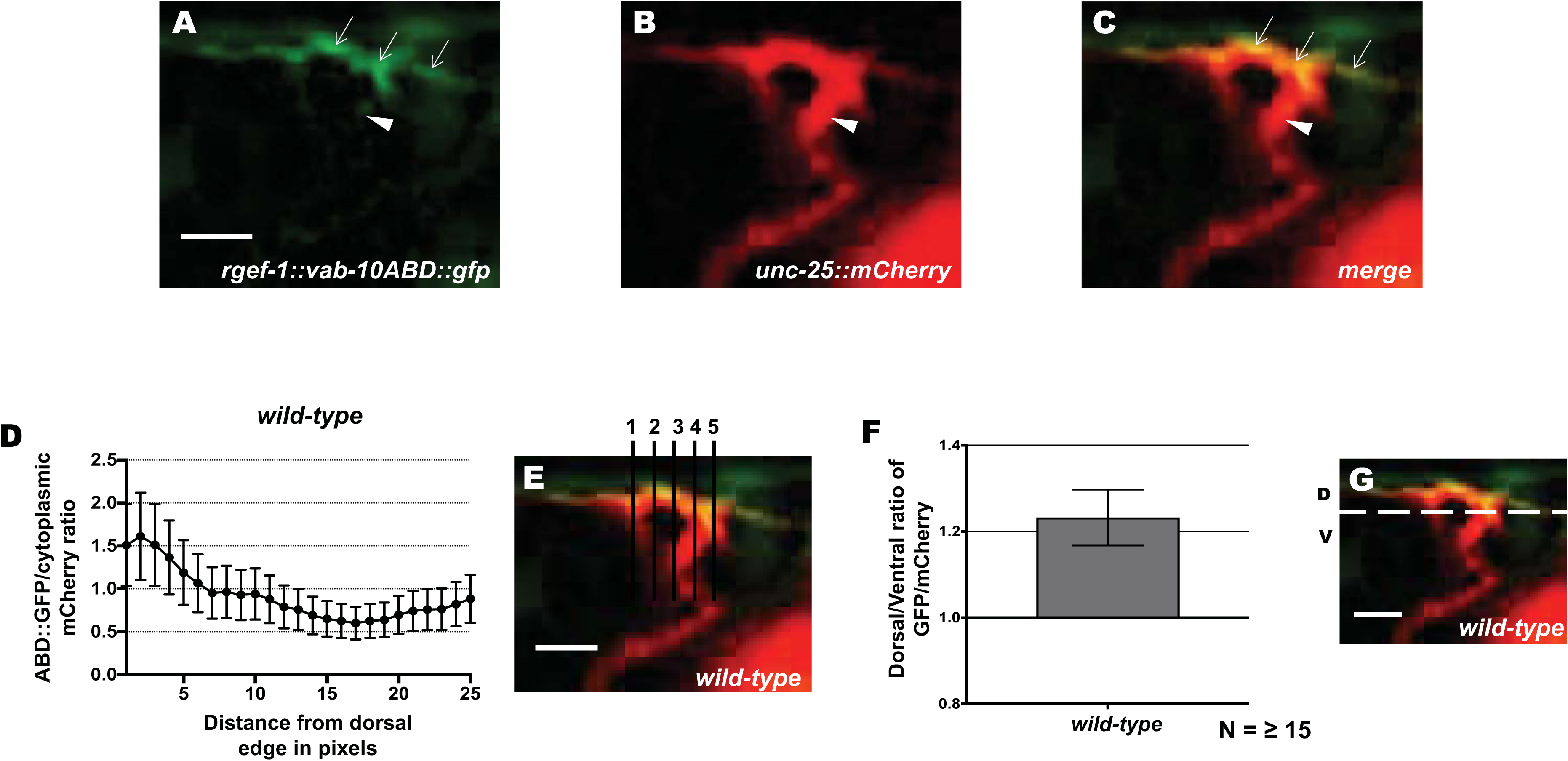
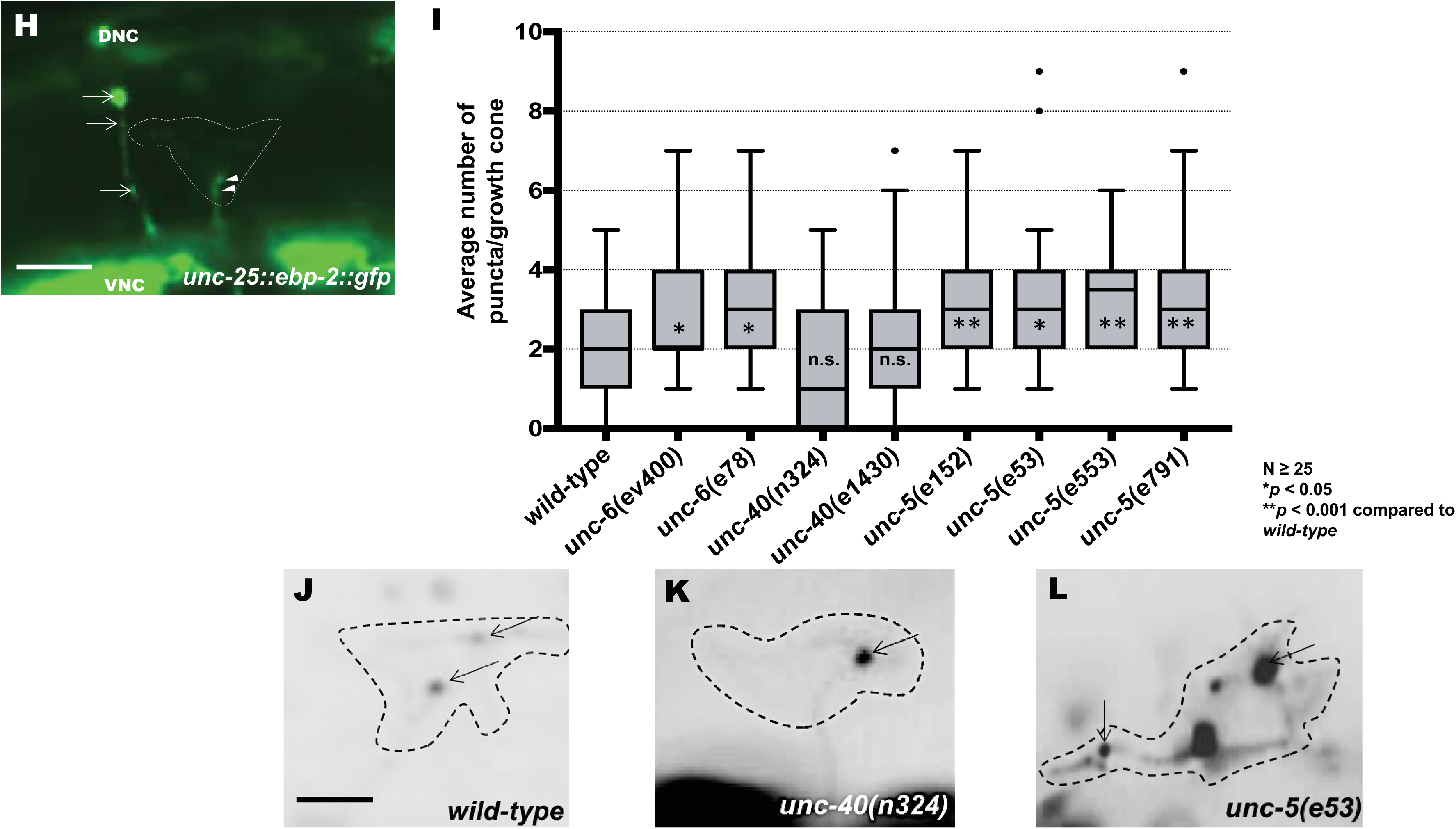
Dorsally-polarized F-actin and EBP-2::GFP accumulation. **(A)** VAB-10ABD::GFP accumulation at the dorsal edge of a wild-type VD growth cone (arrows). Ventral region of the growth cone with little VAB-10ABD::GFP accumulation (arrow heads). **(B)** mCherry growth cone volume marker. **(C)** Merge. Dorsal is up and anterior is left. Scale bar: 5μm in A-C. **(D-G).** A representative line plot of a wild-type VD growth cone as previously described (Norris and Lundquist 2011). **(D)** A graph representing the pixel intensity ratio (arbitrary units) of GFP/mCherry (y-axis) against the distance from the dorsal growth cone edge. **(E)** For each growth cone, five lines were drawn as shown and the pixel intensity ratios were averaged (error bars represent standard deviation). **(F)** The average dorsal-to-ventral ratio of GFP/mCherry in wild-type from multiple growth cones (≥15). Error bars represent the standard error of the mean of the ratios from different growth cones. **(G)** Growth cones were divided into dorsal and ventral halves, and the average intensity ratio of VAB-10ABD::GFP/mCherry was determined for each half and represented in (F). The scale bars in (E) and (G) represent 5μm. **(H)** A wild-type VD growth cone with *Punc-25::ebp-2::*gfp expression from the *lqIs279* transgene. The extent of the growth cone body is highlighted by a dashed line. Arrows point to EBP-2::GFP puncta in the axons of a DD neuron. Arrowheads point to puncta in the VD growth cone. VNC is the ventral nerve cord, and DNC is the dorsal nerve cord. Scale bar: 5μm. **(I)** Box-and-whiskers plot of the number of EBP-2::GFP puncta in the growth cones of different genotypes (≥25 growth cones for each genotype). The grey boxes represent the upper and lower quartiles, and error bars represent the upper and lower extreme values. Dots represent outliers. *p* values were assigned using the two-sided *t*-test with unequal variance. **(J-L)** Growth cones of different genotypes, with EBP-2::GFP puncta indicated with arrows. Dashed lines indicate the growth cone perimeter. Dorsal is up and anterior is left. Scale bar: 5μm.

The MT+-end binding protein EBP-2 fused to GFP has been used previously to monitor MT+ ends in embryos and neuronal processes in *C. elegans* (Srayko *et al.* 2005; Kozlowski *et al.* 2007; Maniar *et al.* 2012; Yan *et al.* 2013; Kurup *et al.* 2015). We expressed *ebp-2::gfp* in the VD/DD neurons using the *unc-25* promoter. Puncta of EBP-2::GFP fluorescence were distributed along the length of commissural axons (arrows in Figure 2H) and in growth cones (arrowheads in Figure 2H and arrows in Figure 2J-L). In wild-type VD growth cones, an average of 2 EBP-2::GFP puncta were observed in the growth cone itself (Figure 2I). These were present at the growth cone base (arrowheads in Figure 2H) as well as in the growth cone periphery (Figure 2J). These data show that in wild-type, EBP-2::GFP puncta were abundant in the axon as previously observed, but relatively rare in the growth cone.

### UNC-5 and UNC-6 affect VD growth cone F-actin dorsal asymmetry and EBP-2::GFP accumulation

In *unc-6* and *unc-5* mutants, VAB-10ABD::GFP dorsal asymmetry in the VD growth cone was abolished (Figure 3A and D and (Norris and Lundquist 2011)). VAB-10ABD::GFP was observed at the growth cone periphery, but often in lateral and even ventral positions (Figure 3D). This loss of VAB-10ABD::GFP asymmetry was accompanied by a corresponding loss of dorsal asymmetry of filopodial protrusion, which occurred all around the growth cone in *unc-5* and *unc-6* mutants (Norris and Lundquist 2011). *unc-40* null mutants displayed relatively normal VD growth cone protrusion compared to wild-type and also showed no effect on VAB-10ABD::GFP distribution (Figure 3A and C and (Norris and Lundquist 2011). These results suggest that UNC-6 and UNC-5 normally control distribution of F-actin to the dorsal leading edge of the VD growth cone, and thus restrict filopodial protrusion to the dorsal leading edge.

**Figure 3.**
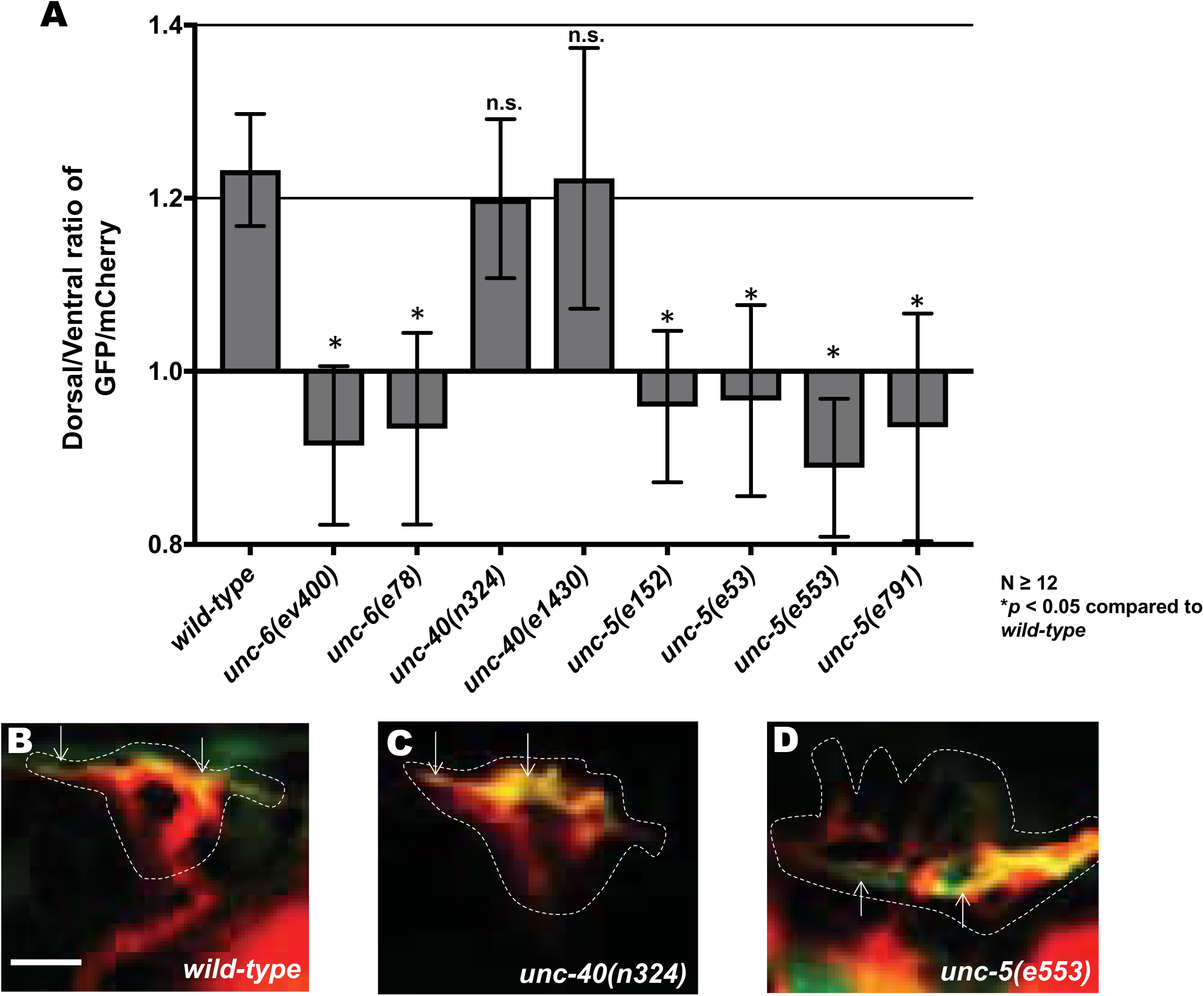
Dorsal F-actin polarity is lost in *unc-5* and *unc-6* mutants. **(A)** The average dorsal-to-ventral ratio of GFP/mCherry from multiple growth cones (≥12) from different genotypes as described in Figure 1. Asterisks (*) indicate the significance of difference between wild-type and the mutant phenotype (\**p* < 0.05) (two-tailed *t*-test with unequal variance between the ratios of multiple growth cones of each genotype). Error bars represent the standard error of the mean **(B-D)** Representative merged images of VD growth cones with cytoplasmic mCherry in red (a volumetric marker) and the VAB-10ABD::GFP in green. Areas of overlap are yellow. Dashed lines indicate the perimeter of the growth cone. Scale bar: 5 μm.

*unc-5* and *unc-6* mutants displayed significantly increased numbers of EBP-2::GFP puncta in VD growth cones and filopodial protrusions (Figure 2I and L). In some mutant growth cones, more than eight puncta were observed, whereas wild-type never showed more than five. *unc-40* mutants displayed no significant increase in EBP-2::GFP puncta accumulation (Figure 2I and K). Sizes of EBP-2::GFP puncta in *C. elegans* neurons were previously found to be on the order of the smaller puncta we observe in wild type (~100nm) (Figure 2J) (Maniar *et al.* 2012). In *unc-5* and *unc-6* mutants, we observed larger puncta (~0.5-1μm) (Figure 2L). We do not understand the nature of the distinct puncta sizes, but the same integrated transgene was used to analyze wild-type and mutants. This suggests that puncta size and number are an effect of the mutant and not transgene variation. In sum, these studies suggest that UNC-5 and UNC-6 are required for the dorsal bias of F-actin accumulation in VD growth cones, and might be required to restrict MT + end entry into VD growth cones as represented by increased numbers of EBP-2::GFP puncta in mutant growth cones.

### EBP-2::GFP puncta accumulation and loss of growth cone F-actin polarity in *unc-5* mutants is not dependent on functional UNC-6 or UNC-40

We found that in *unc-5(e53)*; *unc-40(n324)* and *unc-5(e53); unc-6(ev400)* double mutants, VAB-10ABD::GFP distribution resembled that of *unc-5* mutants alone (i.e. was randomized in the growth cone) (Figure 4A-D). Likewise, EBP-2::GFP accumulation resembled *unc-5*, with increased EBP-2::GFP puncta compared to wild-type or *unc-40* alone (Figure 4E-H). Thus, while UNC-6 and UNC-40 activities were required for the excess growth cone protrusion observed in *unc-5* mutants, they were not required for randomized F-actin or for increased EBP-2::GFP accumulation. These results suggest that UNC-6 and UNC-40 have a role in protrusion that is independent of UNC-5-mediated F-actin dorsal accumulation and EBP-2::GFP accumulation. Consistent with this idea, *unc-6(ev400)* null mutants alone displayed loss of F-actin polarity and increased EBP-2::GFP puncta (Figures 3 and 4), but not increased protrusion (Figure 1) (Norris and Lundquist 2011), suggesting that UNC-6 is required for both UNC-40-mediated protrusion and UNC-5-mediated inhibition of protrusion.

**Figure 4.**
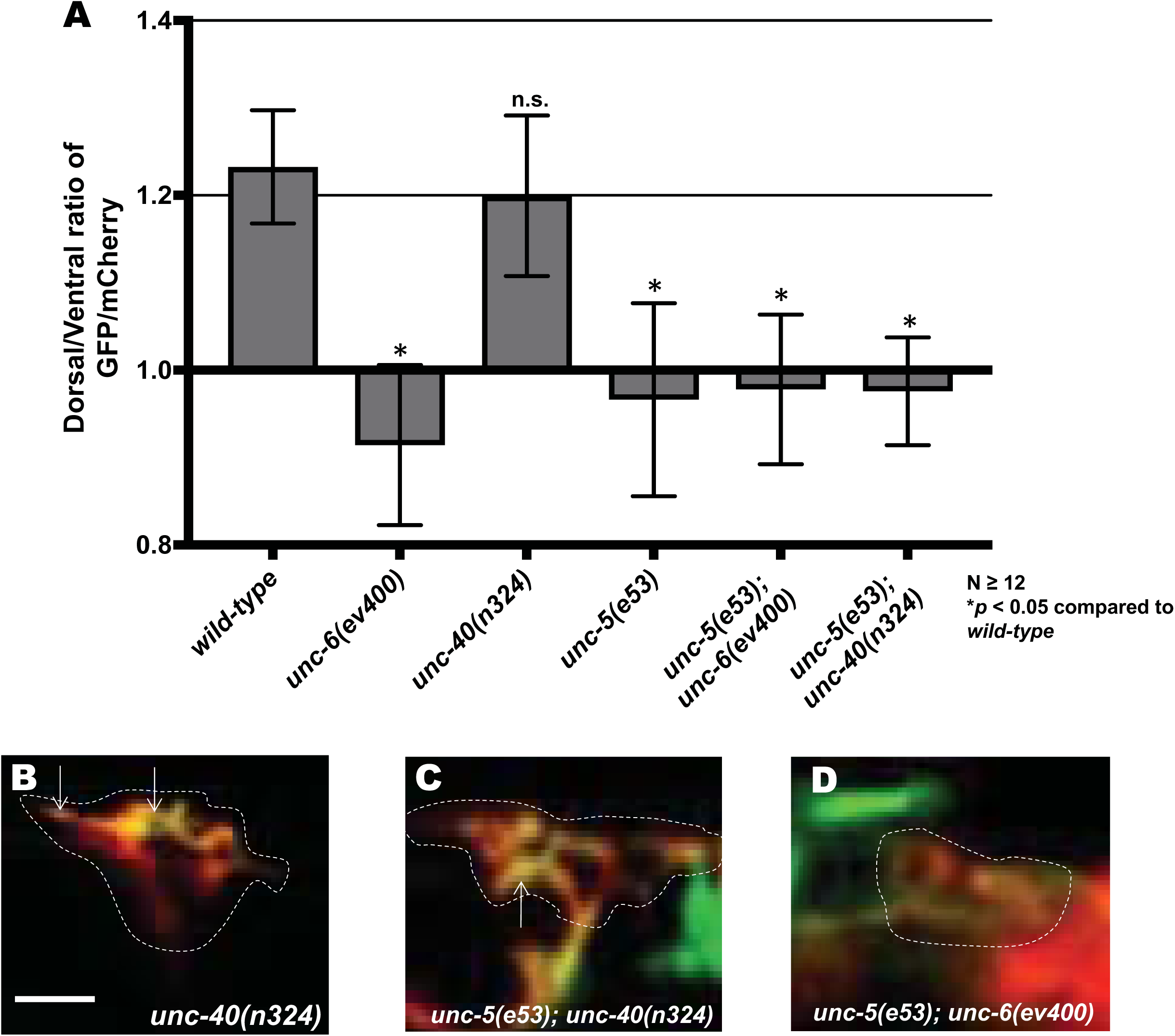
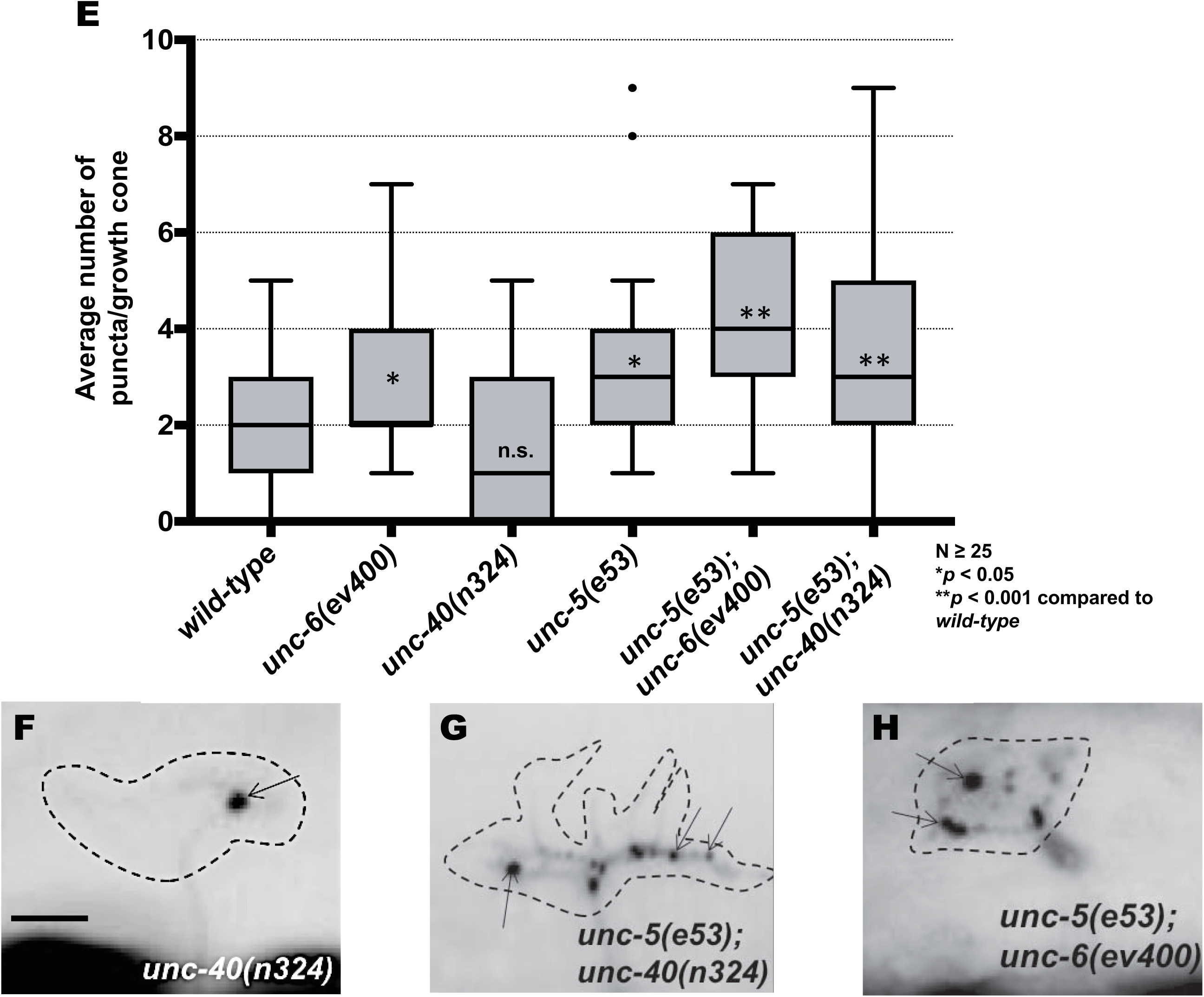
EBP-2::GFP puncta accumulation and loss of growth cone F-actin polarity in *unc-5* mutants is not dependent on functional UNC-6 or UNC-40. **(A)** The average dorsal-to-ventral ratio of GFP/mCherry from multiple growth cones in wild-type and mutant animals as described in Figure 3. **(B-D)** Representative images of VD growth cones with cytoplasmic mCherry in red (a volumetric marker) and the VAB-10ABD::GFP in green as described in Figure 1. Scale bar: 5 μm. **(E)** Quantification of average number of EBP-2::GFP puncta in wild-type and mutant animals as described in Figure 2. **(F-H)** Fluorescence micrographs of EBP-2::GFP expression in VD growth cones. Arrows point to EBP-2::GFP puncta Scale bar: 5μm

### UNC-73 Rac GEF activity controls growth cone F-actin polarity

Previous studies showed that the Rac GTP exchange factor activity of UNC-73 was required to inhibit growth cone protrusion downstream of UNC-5 (Norris *et al.* 2014). *unc-73(rh40),* which specifically eliminates the Rac GEF activity of the molecule (Figure 5A) (Steven *et al.* 1998), resulted in excessive filopodial protrusion (Figure 5B, C, and E, and (Norris *et al.* 2014)). *unc-73(rh40)* mutants displayed a loss of VAB-10ABD::GFP dorsal symmetry in the VD growth cone similar to *unc-5* and *unc-6* mutants (Figure 6A and C). However, *unc-73(rh40)* mutants did not show significantly increased EBP-2::GFP puncta distribution compared to wild-type (Figure 6E and G). Thus, despite having excessively protrusive growth cones, *unc-73(rh40)* mutants did not display increased EBP-2::GFP puncta number. This indicates that the increased numbers of EBP-2::GFP puncta observed in *unc-5* and *unc-6* mutants was not simply due to larger growth cone size. This result also indicates that excess growth cone protrusion can occur in the absence of increased numbers of EBP-2::GFP puncta.

**Figure 5.**
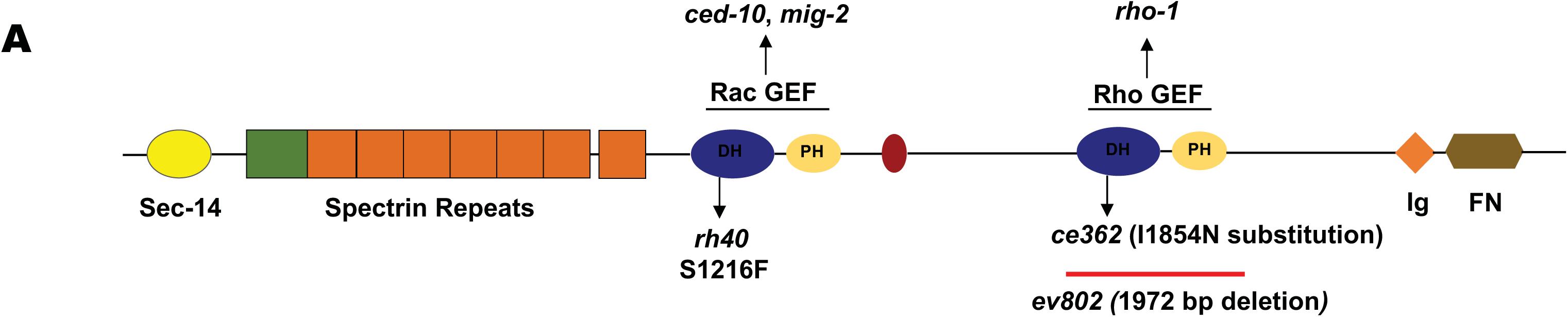
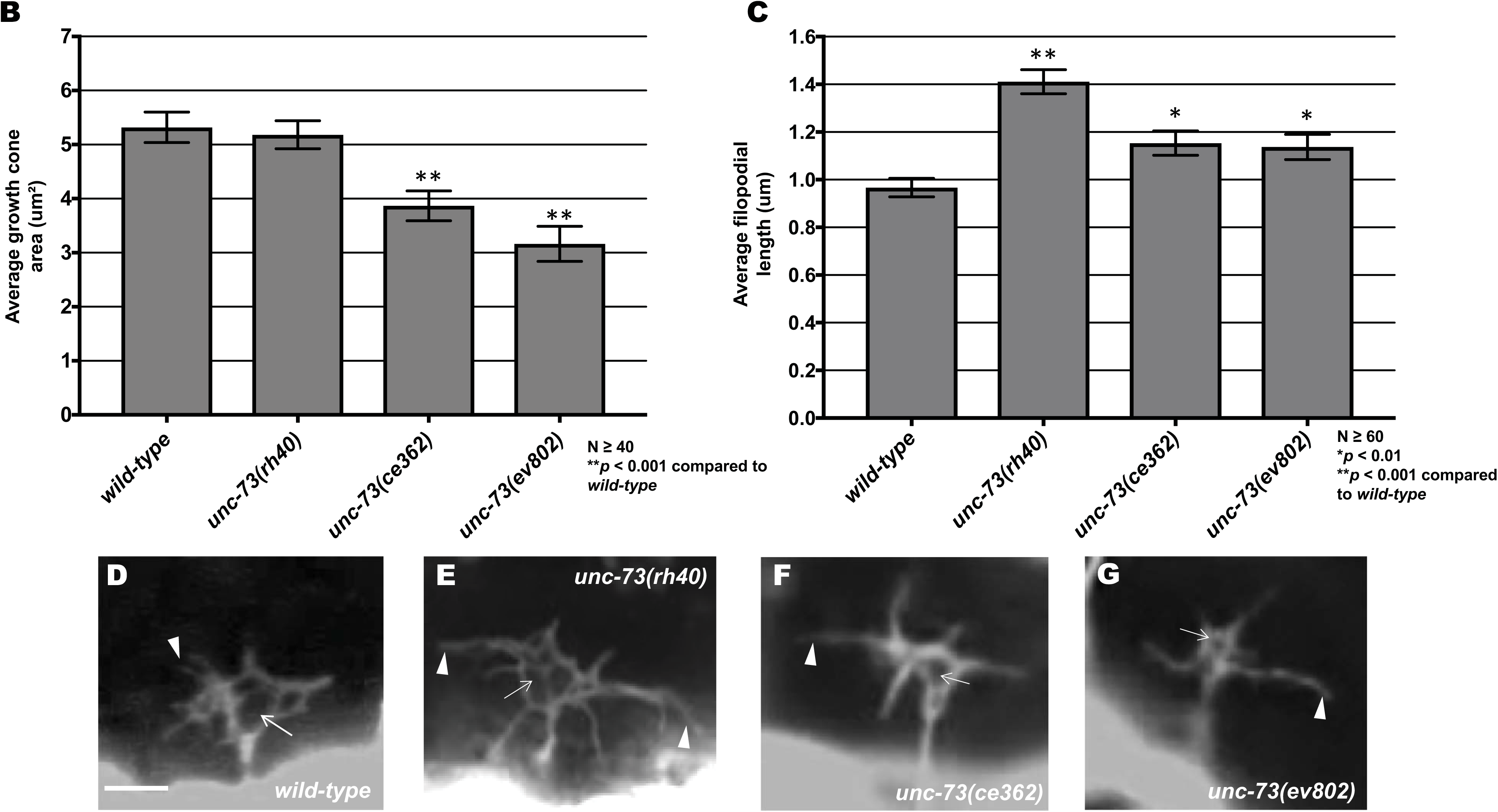
UNC-73/Trio alleles have distinct effects on VD growth cone protrusion. **(A)** A diagram of the full-length UNC-73/Trio molecule. The *rh40, ce362,* and *ev802* mutations are indicated. **(B)** Graphs of the average growth cone area and filopodial length in wild-type and *unc-73* mutants, as described in (Norris and Lundquist 2011) (See Methods). Significance was determined by a two-sided *t*-test with unequal variance. **(D-G)** Fluorescence micrographs of VD growth cones with *Punc-25::gfp* expression from the transgene *juIs76*. Arrows point to the growth cone body, and arrowheads to filopodial protrusions. Scale bar: 5μm.

**Figure 6.**
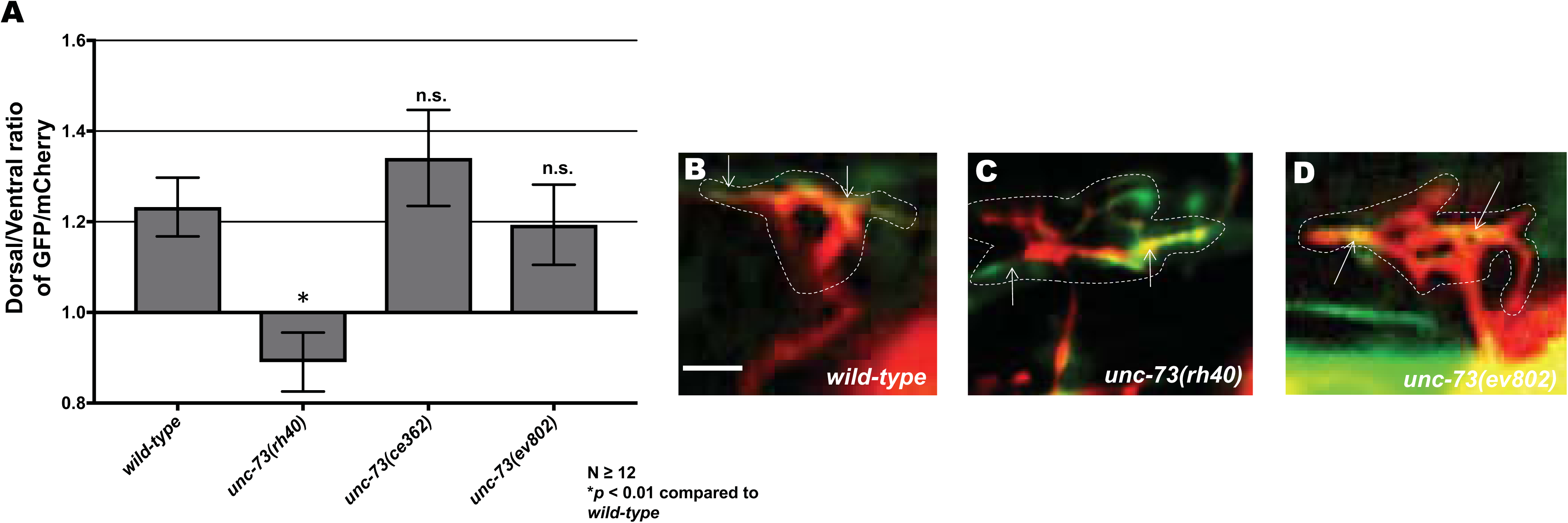
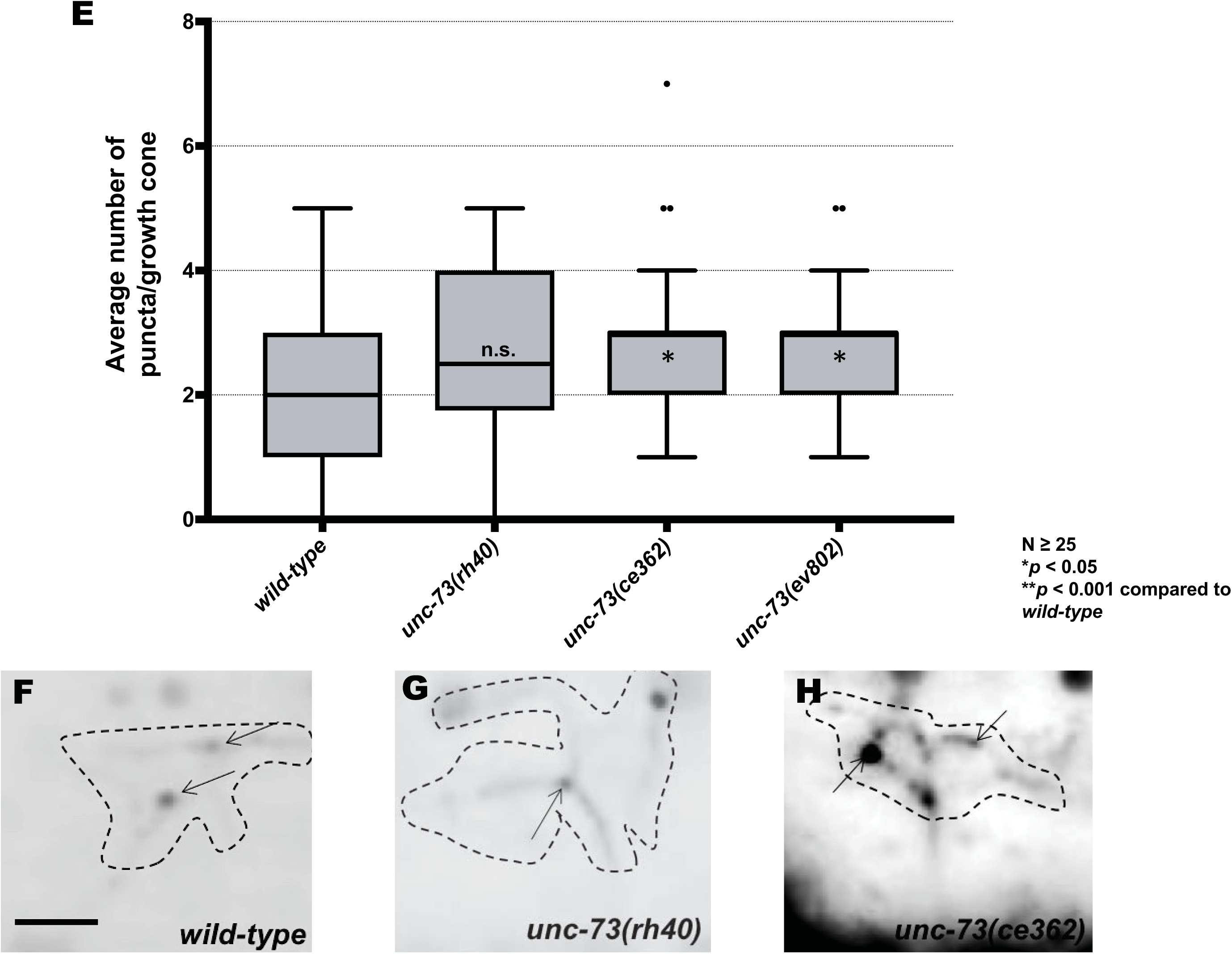
The Rac GEF activity of UNC-73/Trio affects F-actin polarity but not EBP-2::GFP puncta accumulation. **(A)** The average dorsal-to-ventral ratio of GFP/mCherry from multiple growth cones in wild-type and mutant animals as described in Figures 1 and 3. **(C-E)** Representative merged images of VD growth cones with cytoplasmic mCherry in red (a volumetric marker) and the VAB-10ABD::GFP in green, as in Figure 1. Areas of overlap are yellow (arrows). Scale bar: 5 μm. **(F)** Quantification of average number of EBP-2::GFP puncta in wild-type and mutant animals as described in Figure 2. **(G-J)** Fluorescence micrographs of VD growth cones showing EBP-2::GFP puncta (arrows). Scale bar: 5μm.

The C-terminal GEF domain of UNC-73 controls the Rho GTPase (Figure 5A) (Spencer *et al.* 2001) and has been shown to affect motility and normal synaptic neurotransmission (Steven *et al.* 2005; Hu *et al.* 2011). *unc-73(ce362)* is a missense mutation in the Rho GEF domain (Figure 5A) (Williams *et al.* 2007; Hu *et al.* 2011; Mcmullan *et al.* 2012) and *unc-73(ev802)* is a 1,972 bp deletion which completely deletes the Rho GEF domain (Figure 5A) (Williams *et al.* 2007). *unc-73(ce362)* and *unc-73(ev802)* displayed reduced growth cone body size, and increased filopodial length (Figure 5B, C, F, and G) Neither *unc-73(ce362)* nor *unc-73(ev802)* had an effect on growth cone F-actin dorsal accumulation (Figure 6A and D), but both showed a significant increase in growth cone EBP-2::GFP puncta (Figure 6E and 6H). Thus, Rho GEF activity of UNC-73/Trio might have distinct effects on growth cone morphology compared to Rac GEF activity.

### The Rac GTPases CED-10 and MIG-2 affect F-actin polarity but not EBP-2::GFP puncta accumulation in VD growth cones

The Rac GTPases CED-10/Rac and MIG-2/RhoG have been shown to redundantly control axon guidance (Lundquist *et al.* 2001; Struckhoff and Lundquist 2003). CED-10 and MIG-2 act with UNC-40 to stimulate protrusion in axons attracted to UNC-6/Netrin (Demarco *et al.* 2012), and to inhibit growth cone protrusion with UNC-5-UNC-40 in the repelled VD growth cones (Norris *et al.* 2014). The Rac GEF TIAM-1 acts with CED-10 and MIG-2 to stimulate protrusion (Demarco *et al.* 2012), and the Rac GEF UNC-73/Trio acts in the anti-protrusive pathway (Norris *et al.* 2014).

The VD growth cones of *mig-2; ced-10* double mutants resembled wild-type, except that the filopodial protrusions had a longer maximal length and were longer lasting (Norris *et al.* 2014). This subtle phenotype might represent the fact that the molecules have roles in both pro- and anti-protrusive pathways. *mig-2(mu28)* and *ced-10(n1993)* single mutants and *ced-10; mig-2* double mutants all showed significant F-actin polarity defects (Figure 7A-D), consistent with the idea that the GEF domain of Trio affects actin organization through Rac activation. However, VD/DD axon guidance defects in *mig-2* and *ced-10* single mutants were much less severe than the *mig-2; ced-10* double mutants and *unc-73(rh40)* (Norris *et al.* 2014). *unc-73(rh40)* and *mig-2; ced-10* double mutants might have additional effects on axon guidance compared to *mig-2* and *ced-10* double mutants not observed in these studies. Similar to *unc-5; unc-40* double mutants, loss of asymmetry of F-actin did not result in excess protrusion in *mig-2* and *ced-10* single mutants.

**Figure 7.**
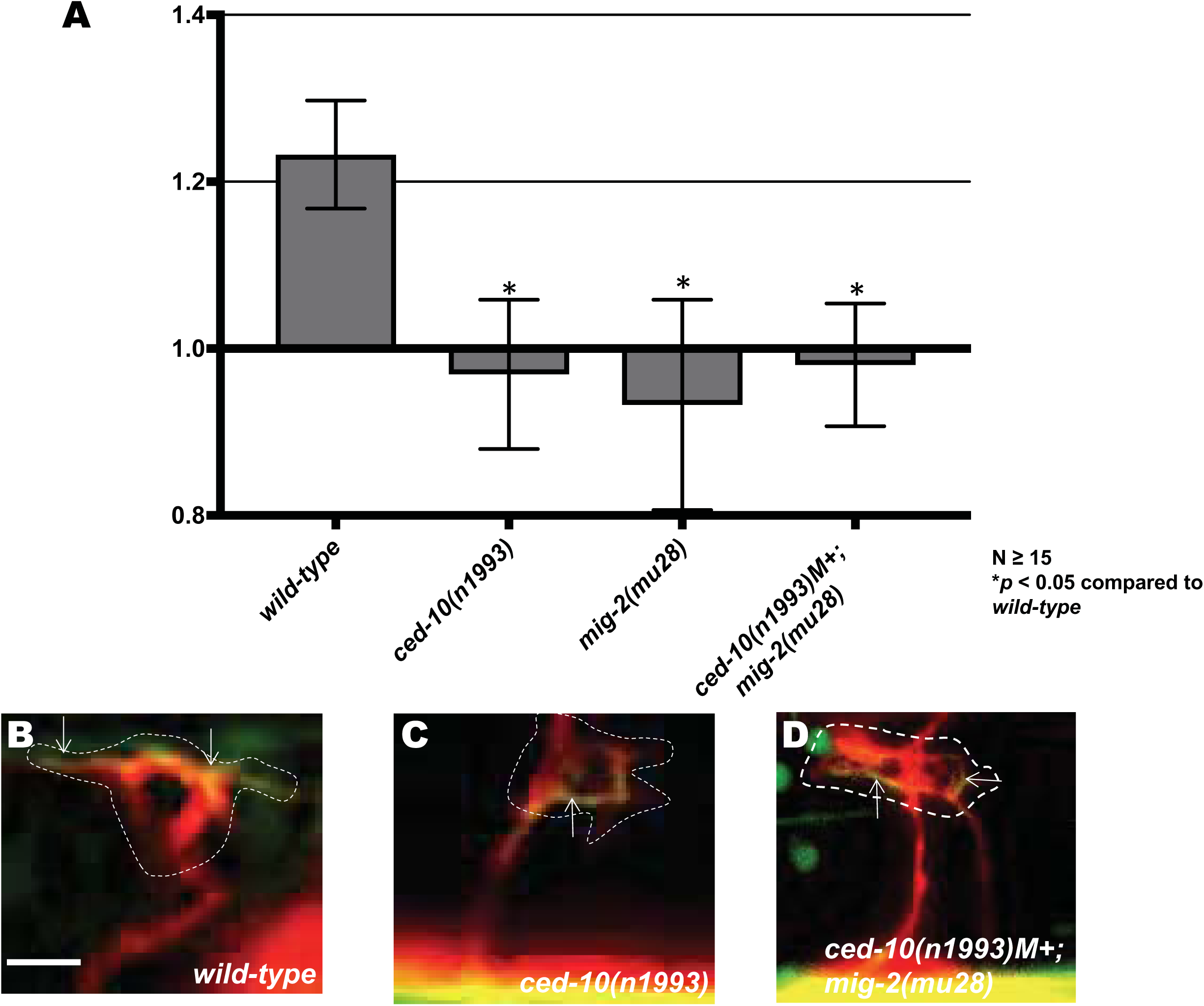
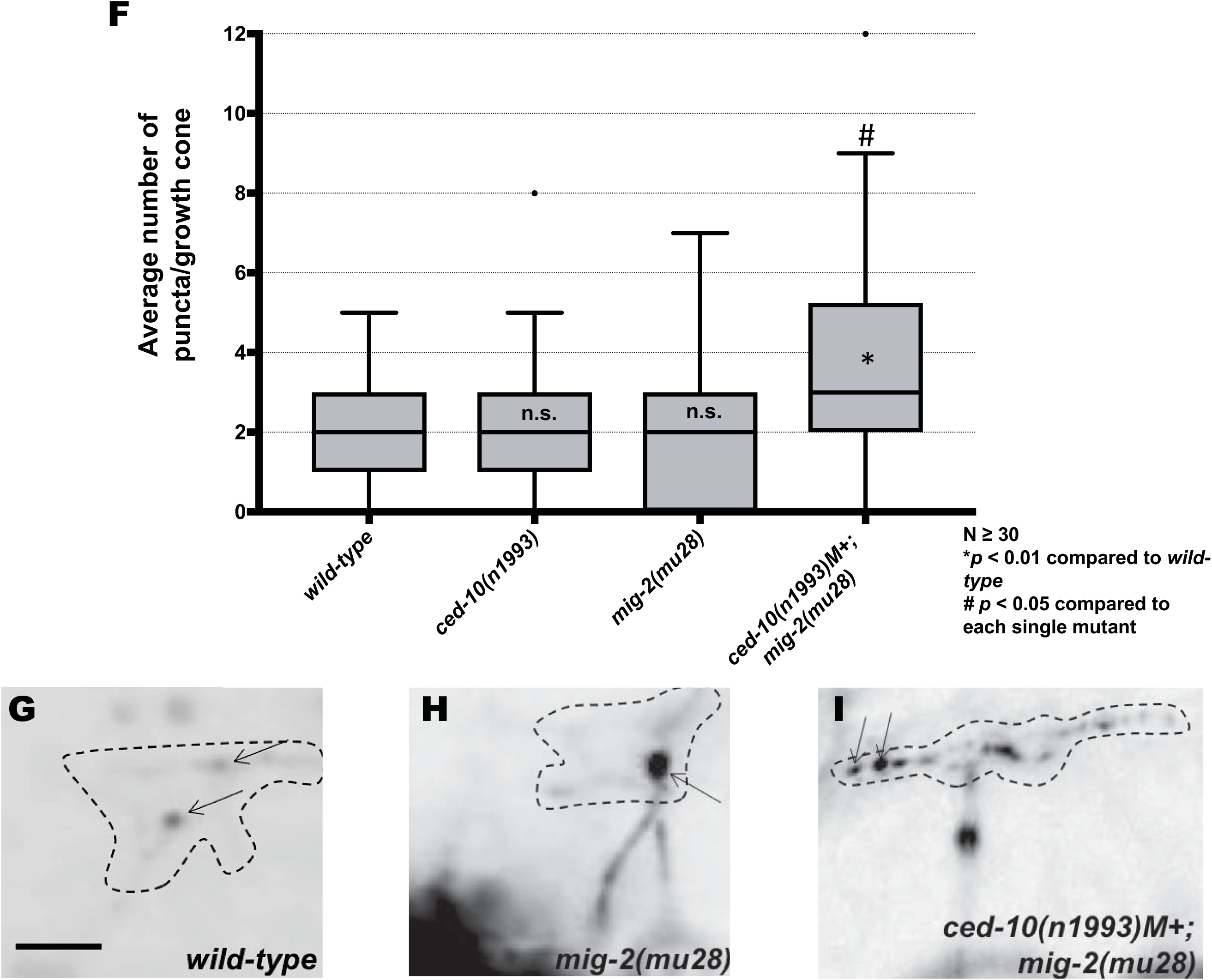
The Rac GTPases CED-10 and MIG-2 individually affect F-actin polarity and are redundant for EBP-2::GFP puncta accumulation. **(A)** The average dorsal-to-ventral ratio of GFP/mCherry from multiple growth cones in wild-type and mutant animals as described in Figures 1 and 3. **(B-D)** Representative merged images of VD growth cones with cytoplasmic mCherry in red (a volumetric marker) and the VAB-10ABD::GFP in green as in Figure 1. Scale bar: 5 μm. **(E)** Quantification of average number of EBP-2::GFP puncta in wild-type and mutant animals as described in Figure 2. **(F-H)** Fluorescence micrographs of VD growth cones with EBP-2::GFP puncta indicated arrows. Scale bar: 5μm.

*ced-10* and *mig-2* single mutants had no significant effect on EBP-2 distribution in the VD growth cone (Figure 7E and H). However, *ced-10; mig-2* double mutants showed a significant increase in EBP-2 puncta distribution in the VD growth cone and filopodial protrusions as compared to wild-type and the single mutants alone (Figure 7E and I). This result suggests that the Rac GTPases CED-10 and MIG-2 act redundantly in limiting EBP-2 puncta distribution in the VD growth cone. This also indicates that MIG-2 and CED-10 have a role in limiting EBP-2::GFP puncta that is independent of UNC-73/Trio Rac GEF activity.

### UNC-33/CRMP and UNC-44/ankyrin are required for F-actin polarity and restricting EBP-2::GFP from the VD growth cone

Previous studies showed that UNC-33/CRMP and UNC-44/ankyrin act downstream of UNC-5 and Rac GTPases to limit growth cone protrusion (Norris *et al.* 2014). *unc-33* and *unc-44* mutants randomized F-actin polarity similar to *unc-5* and *unc-73(rh40)* (Figure 8A-D). *unc-33* and *unc-44* also displayed a significant increase in EBP-2::GFP puncta in the growth cone and protrusions (Figure 8E-H). Thus, UNC-33 and UNC-44 are both required for dorsal F-actin asymmetry as well as restriction of EBP-2::GFP growth cone puncta. That *unc-33* and *unc-44* phenotypes are similar to *unc-5* is consistent with the previous genetic interactions placing UNC-33 and UNC-44 in the UNC-5 pathway.

**Figure 8.**
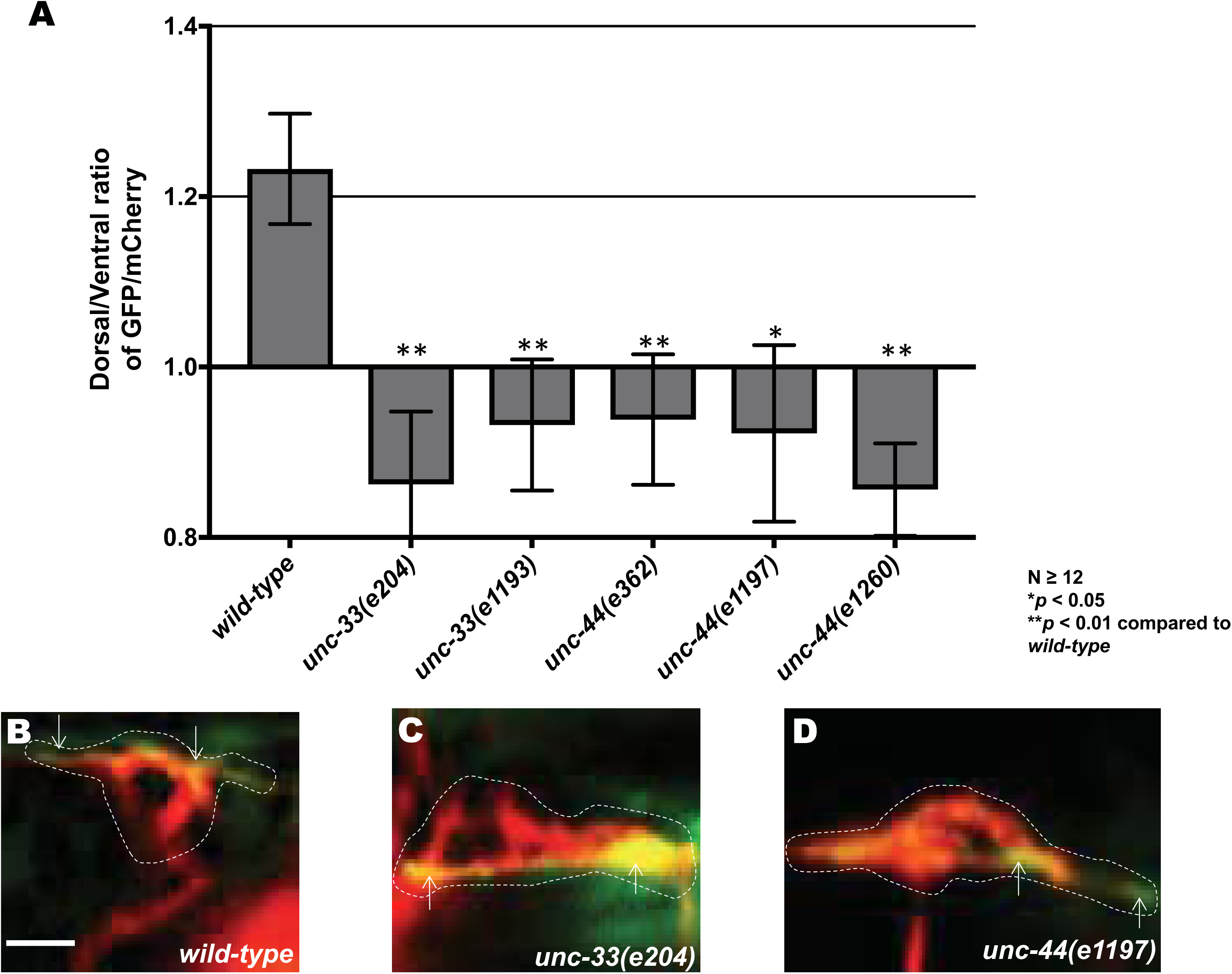
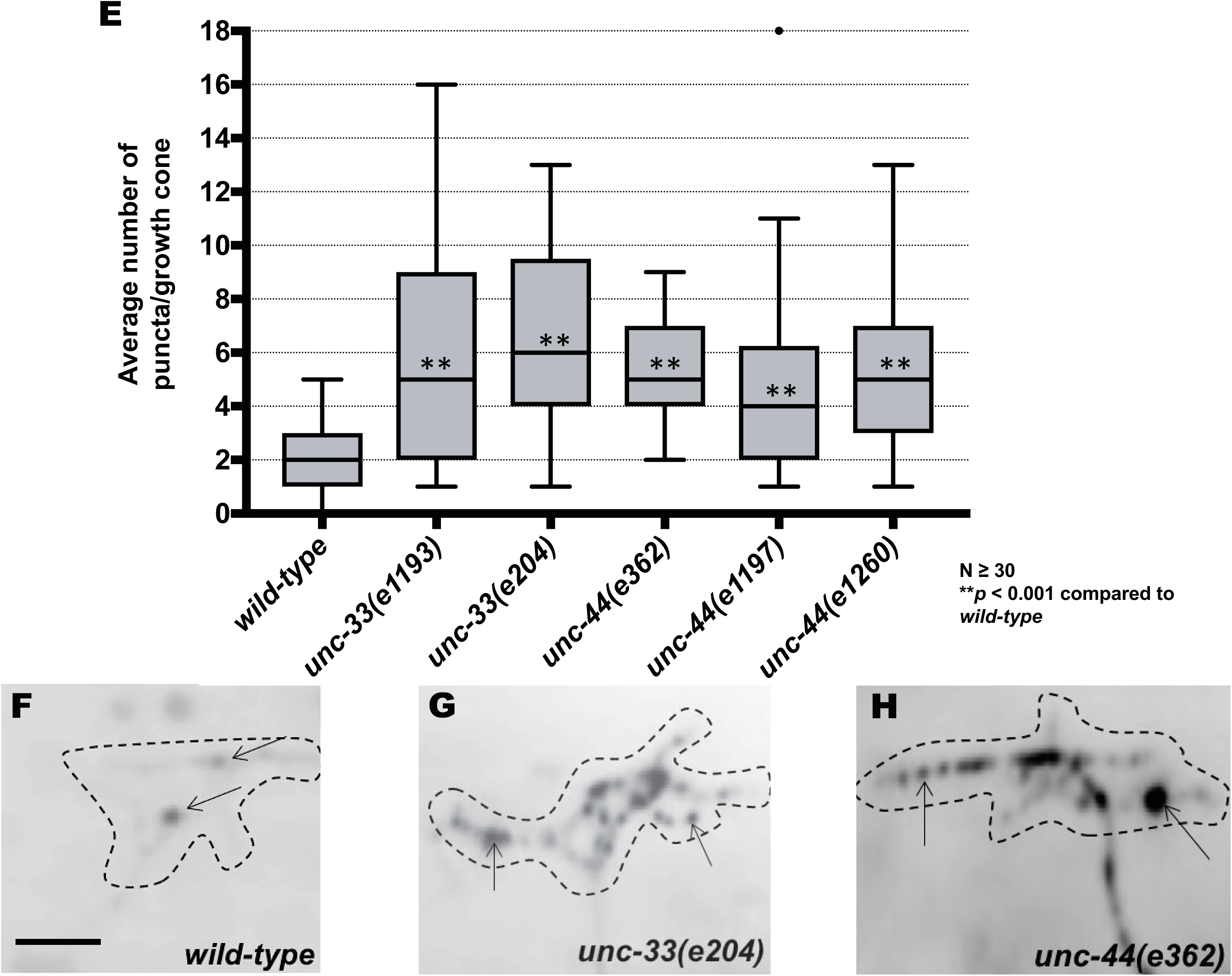
*unc-33* and *unc-44* mutants disrupt F-actin polarity and affect EBP-2::GFP puncta accumulation. **(A)** The average dorsal-to-ventral ratio of GFP/mCherry from multiple growth cones in wild-type and mutant animals as described in Figure 3. **(B-D)** Representative merged images of VD growth cones with cytoplasmic mCherry in red (a volumetric marker) and the VAB-10ABD::GFP in green, as in Figure 1. Scale bar: 5 μm. **(E)** Quantification of average number of EBP-2::GFP puncta in wild-type and mutant animals as in Figure 2. **(F-H)** Fluorescence micrographs of VD growth cones with EBP-2::GFP puncta indicate by arrows. Scale bar: 5μm.

### Constitutive activation of UNC-40, UNC-5, CED-10 and MIG-2 affects F-actin polarity and EBP-2 distribution

The heterodimeric receptor UNC-5-UNC-40 is required for inhibition of growth cone protrusion in UNC-6/netrin repulsive axon guidance (Norris and Lundquist 2011; Norris *et al.* 2014). Constitutive activation of UNC-40 and UNC-5 by addition of an N-terminal myristoylation signal to their cytoplasmic domain (Gitai *et al.* 2003; Norris and Lundquist 2011) causes a significant decrease in VD growth cone protrusiveness, with a reduction in growth cone area and filopodial protrusions (Demarco *et al.* 2012; Norris *et al.* 2014). The Rac GTPases CED-10 and MIG-2 have been shown to act in both stimulation and inhibition of growth cone protrusion (Demarco *et al.* 2012; Norris *et al.* 2014). The constitutively activated Rac GTPases CED-10(G12V) and MIG-2(G16V) also cause an inhibited VD growth cone phenotype similar to *myr::unc-40* and *myr::unc-5* (Norris et al. 2014).

We assayed VAB-10ABD::GFP and EBP-2::GFP distribution in the VD growth cones of these various activated molecules. All four (MYR::UNC-5, MYR::UNC-40, CED-10(G12V), and MIG-2(G16V) showed peripheral accumulation around the entire growth cone, with the dorsal bias lost (Figure 9A-D). Furthermore, we observed significantly fewer EBP-2::GFP puncta compared to wild type in each case (Figure 9E-H). Thus, constitutive activation of UNC-5, UNC-40, CED-10, and MIG-2 might lead to F-actin accumulation around the entire periphery of the growth cone, as opposed to dorsal bias, and might restrict MT + ends from accumulating in the growth cone.

**Figure 9.**
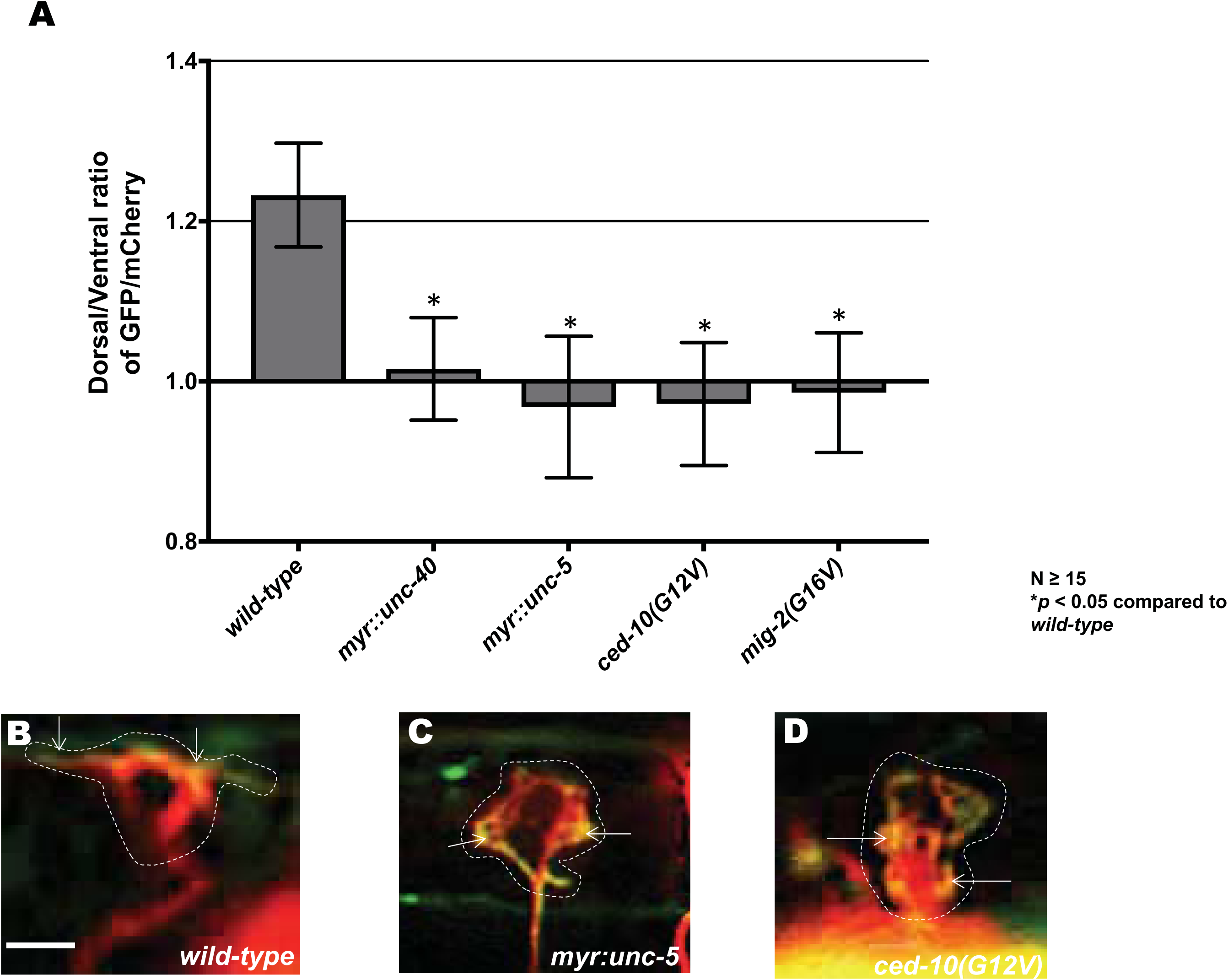
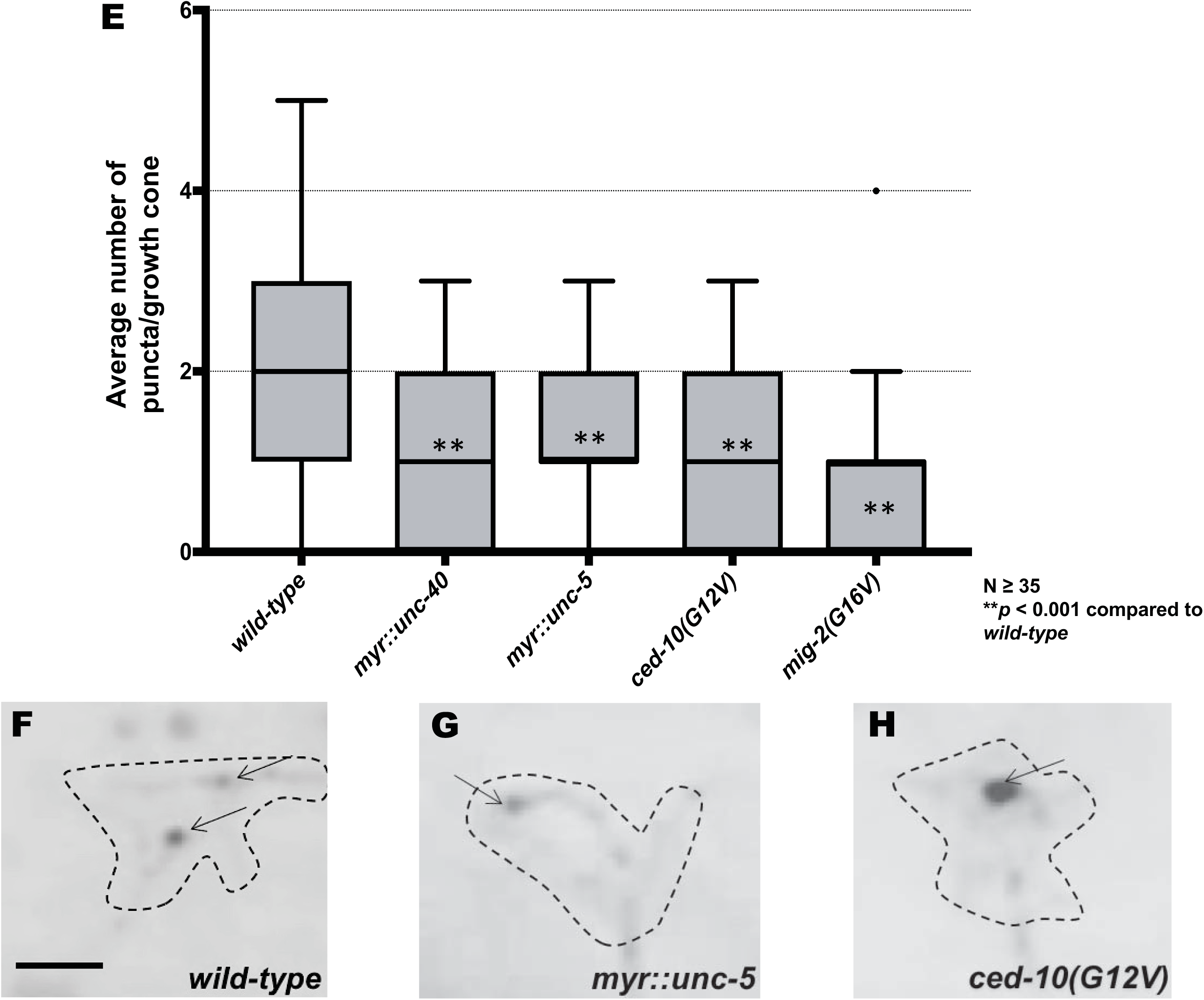
Constitutive activation of UNC-40, UNC-5, CED-10 and MIG-2 affects F-actin polarity and EBP-2 distribution. **(A)** The average dorsal-to-ventral ratio of GFP/mCherry from multiple growth cones in wild-type and mutant animals as described in Figure 3. **(B-D)** Representative merged images of VD growth cones with cytoplasmic mCherry in red (a volumetric marker) and the VAB-10ABD::GFP in green. Scale bar: 5 μm. **(E)** Quantification of average number of EBP-2::GFP puncta in wild-type and mutant animals as described in Figure 2. **(F-H)** Fluorescence micrographs of VD growth cones with EBP-2::GFP puncta indicated by arrows. Scale bar: 5μm.

## Discussion

Netrins are thought to regulate dorsal-ventral axon guidance through a conserved mechanism involving ventral expression of Netrin coupled with expression of UNC-40/DCC receptors on attracted axons and UNC-5 receptors on repelled axons. Recent studies have highlighted the previously-underappreciated complexity of UNC-6/Netrin function in axon guidance. In *C. elegans,* UNC-5 acts in axons that grow toward Netrin to focus UNC-40 activity (Levy-Strumpf and Culotti 2014; Yang *et al.* 2014; Limerick *et al.* 2018), and both UNC-40 and UNC-5 act in the same growth cone to mediate protrusion in directed guidance away from UNC-6/Netrin (Norris and Lundquist 2011; Norris *et al.* 2014). In this work, we analyze three aspects of VD growth cone morphology (growth cone protrusion, F-actin accumulation, and EBP-2::GFP accumulation) to probe the roles of UNC-6/Netrin, UNC-40/DCC, and UNC-5 in growth away from UNC-6/Netrin. We find that UNC-6/Netrin signaling coordinates these growth cone features to result in directed growth away from it.

Previous studies suggested that UNC-6/Netrin and the receptor UNC-5-UNC-40 inhibit growth cone protrusion and are required to polarize F-actin to the dorsal protrusive leading edge in repelled VD growth cones in *C. elegans* (Norris and Lundquist 2011; Norris *et al.* 2014). Furthermore, previous studies defined a new signaling pathway downstream of UNC-5-UNC-40 in repulsive axon guidance and inhibition of growth cone protrusion. This pathway consists of the Rac GTP exchange factor UNC-73/Trio, the Rac GTPases CED-10 and MIG-2, and the cytoskeletal interacting molecules UNC-44/ankyrin and UNC-33/CRMP (Norris *et al.* 2014). Loss of function in these molecules led to excess growth cone protrusion, and activation led to constitutively-inhibited growth cone protrusion. Using VAB-10ABD::GFP to visualize F-actin and EBP-2::GFP to visualize MT + ends, we endeavored in this work to define the effects of members of this pathway on the cytoskeleton of the VD growth cone. We found that mutations in *unc-5, unc-6, unc-33,* and *unc-44* led to excessively protrusive growth cones with randomized F-actin distribution and increased numbers of EBP-2::GFP puncta in the growth cones. *unc-40* mutation suppressed the excess protrusion of *unc-5* mutants but not the F-actin randomization nor the excess EBP-2::GFP puncta, suggesting that UNC-40 might have a stimulatory role in protrusion independent of UNC-5. We found that Rac GEF activity was required to inhibit protrusion and for F-actin dorsal bias, but was not required to restrict EBP-2::GFP puncta, suggesting a mechanism distinct from EBP-2::GFP puncta increase that stimulates growth cone protrusion. Finally, we found a complex involvement of the Rac GTPases in protrusion, F-actin bias, and EBP-2::GFP puncta that is consistent with these molecules having both pro- and anti-protrusive roles in the growth cone.

### UNC-5 and UNC-6/Netrin might inhibit MT+- end accumulation in the VD growth cone

Loss of *unc-5* and *unc-6* caused significant mislocalization of F-actin and significantly increased the average number of EBP-2::GFP puncta distribution in VD growth cones (Figures 2 and 3), which have been used to track MT + ends in *C. elegans* neurons and embryos (Srayko *et al.* 2005; Kozlowski *et al.* 2007; Maniar *et al.* 2012; Yan *et al.* 2013; Kurup *et al.* 2015). UNC-6 and UNC-5 might inhibit growth cone protrusion by preventing F-actin formation in the ventral/lateral regions of the growth cone to restrict protrusion to the dorsal leading edge, and by preventing accumulation of MT + ends in the growth cone. In cultured growth cones, MTs are involved in both DCC and UNC5C-mediated axon outgrowth, and DCC and UNC5C physically associate with MTs in a Netrin-dependent manner in cultured cells (Qu *et al.* 2013; Shao *et al.* 2017; Huang *et al.* 2018). Our results suggest a link between UNC-6/Netrin signaling and VD growth cone MTs *in vivo*.

### MTs in the growth cone might be pro-protrusive

Our data show a correlation between MT + ends in growth cones and excess growth cone protrusion. This is consistent with *in vitro* studies of growth cones in which MT + end entry into the growth cone is tightly regulated, is intimately associated with F-actin, and is essential for protrusion and outgrowth (Lowery and Van VACTOR 2009; Dent *et al.* 2011; Vitriol and Zheng 2012; Coles and Bradke 2015). Possibly, MT entry into growth cones serves as a conduit for transport of vesicles, organelles, and pro-protrusive factors involved in actin polymerization that drive filopodial protrusion in *C. elegans,* such as Arp2/3, UNC-115/abLIM, and UNC-34/Enabled (Shakir *et al.* 2006; Shakir *et al.* 2008; Norris *et al.* 2009). This is consistent with results from cultured growth cones showing that MT stabilization results in growth cone turning in the direction of stabilization, and MT destabilization results in growth cone turning away from MT destabilization (Buck and Zheng 2002). Also, MTs are involved in both DCC and UNC5C-mediated axon guidance (Qu *et al.* 2013; Shao *et al.* 2017; Huang *et al.* 2018), and physically associate with UNC5C, which is decoupled by Netrin and associated with growth away from Netrin (Shao *et al.* 2017). We have no evidence that UNC-5 or UNC-40 physically associate with MTs, but these data are consistent with MTs having a pro-protrusive role (i.e. protrusion depends on UNC-40 and MTs, and inhibiting protrusion via UNC-5 results in fewer growth cone MTs).

### UNC-40 might have a pro-protrusive role downstream of EBP-2::GFP puncta and F-actin polarity

*unc-40* alone showed wild-type levels of protrusion (Norris and Lundquist 2011), and here we find that *unc-40* did not affect F-actin polarity or EBP-2::GFP puncta accumulation. Previous work showed that a functional UNC-40 is required for the large protrusive growth cones seen in *unc-5* single mutants (Norris and Lundquist 2011).

However, *unc-5; unc-40* double mutants, despite have smaller growth cones, showed loss of F-actin polarity and excess EBP-2::GFP puncta similar to *unc-5* alone (Figure 4). Thus, UNC-40 might have a role in protrusion that is downstream of F-actin polarity and EBP-2::GFP puncta. In migrating embryonic cells and anchor cell invasion, UNC-40 affects over all F-actin levels but not F-actin polarity (Bernadskaya *et al.* 2012; Wang *et al.* 2014).

Something similar might be occurring in the growth cone, where UNC-40 has a role in actin polymerization but not polarity, which might be determined by UNC-5 or the UNC-5-UNC-40 heterodimer. This is similar to recent results in neurons with axons that grow ventrally toward the UNC-6/Netrin source, which require UNC-5 for ventral guidance (Kulkarni *et al.* 2013; Levy-Strumpf and Culotti 2014). In this case, UNC-40 drives protrusion and is polarized ventrally in the cell body by UNC-6/Netrin. UNC-5 further refines this UNC-40 localization of protrusion and prevents lateral and ectopic protrusions (Kulkarni *et al.* 2013; Yang *et al.* 2014; Limerick *et al.* 2018). Our results suggest that F-actin and EBP-2::GFP accumulation, controlled by UNC-5, are pro-protrusive, and that UNC-40 might act downstream of these events to drive growth cone protrusion.

### The Rac GEF domain of UNC-73 inhibits protrusion independently of restricting MT + ends

Rac GTPases CED-10 and MIG-2 and the UNC-73/Trio Rac GEF have been shown to play central roles in axon guidance (Steven *et al.* 1998; Lundquist *et al.* 2001; Lundquist 2003; STRUCKHOFF AND Lundquist 2003). Rac GTPases CED-10 and MIG-2 are required to both stimulate and inhibit protrusion, with distinct GEFs regulate each of these activities.

TIAM-1 stimulates protrusion (Demarco *et al.* 2012), and UNC-73 limiting protrusion (Norris *et al.* 2014). The *unc-73(rh40)* mutation eliminates the Rac GEF activity of UNC-73 but does not affect Rho GEF activity (Steven *et al.* 1998). *unc-73(rh40)* mutants displayed F-actin polarity defects (Figure 6) consistent with the idea that UNC-73 regulates actin dynamics during cell growth and growth cone migrations (Steven *et al.* 1998; Bateman *et al.* 2000; Lundquist *et al.* 2001; Wu *et al.* 2002). We found that *unc-73(rh40)* had no effect on EBP-2::GFP accumulation in VD growth cones despite having larger, more protrusive growth cones (Figure 6). Despite the large, overly-protrusive growth cones, *unc-73(rh40)* mutants did not display excess EBP-2::GFP puncta as observed in *unc-6, unc-5*, and *unc-33* mutants. This indicates that the excess EBP-2::GFP puncta in *unc-5, unc-6,* and *unc-33* mutants are not due increased growth cone size and protrusion. Furthermore, this suggests that the UNC-73 Rac GEF activity might inhibit protrusion by a mechanism distinct from restricting MT + end entry, possibly by affecting actin polymerization directly. Such a mechanism could involve the flavin monooxygenase (FMOs) FMO-1 and FMO-5, which were recently shown to act downstream of UNC-5 and activated Rac GTPases to inhibit VD growth cone protrusion (Gujar *et al.* 2017). In *Drosophila*, the FMO-containing MICAL molecule causes actin depolymerization by directly oxidizing actin (Hung *et al.* 2010; Hung *et al.* 2011). The *C. elegans* genome does not encode a single MICAL-like molecule containing an FMO plus additional functional domains. In *C. elegans* FMOs might play an analogous role to MICAL in actin regulation and growth cone inhibition.

Mutations in the Rho-specific GEF domain of *unc-73* led to a complex phenotype. Growth cones were slightly smaller with slightly increased filopodial length. F-actin polarity was unaffected, but excess EBP-2::GFP puncta were observed. This phenotype could reflect the role of RHO-1 in the growth cone, or could reflect that these mutations are not specific to the Rho GEF domain and might affect overall function of the molecule. In any event, these mutations display a distinct phenotype compared to *unc-73(rh40)*, which is specific to the Rac GEF activity of UNC-73.

### The Rac GTPases CED-10 and MIG-2 affect F-actin polarity and EBP-2::GFP accumulation

The Rac GEF activity of UNC-73/Trio was required for F-actin polarity but not EBP-2::GFP restriction. However, the *mig-2; ced-10* Rac double mutant displayed both F-actin polarity defects and excess EBP-2::GFP puncta (Figure 7), suggesting that Rac GTPases have an UNC-73/Trio Rac GEF activity-independent role in EBP-2::GFP restriction and thus possibly MT + end restriction from the growth come. Possibly another Rac GEF regulates MIG-2 and CED-10 in MT + end restriction. Despite unpolarized F-actin and excess MT + ends, the growth cones in *mig-2; ced-10* double mutants have only subtly-increased filopodial protrusions, much weaker than *unc-73(rh40)*. This might be due to MIG-2 and CED-10 being required in both pro- and anti-protrusive activities, resulting in an intermediate effect on growth cone protrusion in the double mutant.

*ced-10* and *mig-2* single mutants displayed F-actin polarity defects alone, but did not display excess EBP-2::GFP puncta accumulation. Thus, CED-10 and MIG-2 are individually required for F-actin polarity and act redundantly in MT + end restriction. Despite F-actin polarity defects, protrusion of the *ced-10* and *mig-2* growth cones resembles wild-type. This could again be explained by their roles in both pro- and anti-protrusive activities.

### UNC-33/CRMP and UNC-44/Ankyrin are required to exclude MT+-ends from the VD growth cone

Our previous work showed that the collapsin-response-mediating protein UNC-33/CRMP and UNC-44/ankyrin are required for inhibition of protrusion by UNC-5-UNC-40 and Rac GTPases. Here we show that UNC-33 and UNC-44, similar to UNC-5, are required for F-actin polarity and to restrict MT + end accumulation in the growth cone. CRMPs were first identified as molecules required for growth cone collapse induced by semaphorin-3A through Plexin-A and Neuropilin-1 receptors (Goshima *et al.* 1995; Takahashi *et al.* 1999). CRMP4 knockdown in cultured mammalian neurons led to increased filopodial protrusion and axon branching (Alabed *et al.* 2007), consistent with our findings of UNC-33/CRMP as an inhibitor of protrusion. However, hippocampal neurons from a CRMP4 knock-out mouse exhibited decreased axon extension and growth cone size (Khazaei *et al.* 2014).

CRMPs have various roles in actin and MT organization and function (Khazaei *et al.* 2014). CRMP2 promotes microtubule assembly *in vitro* by interacting with tubulin heterodimers and microtubules to regulate axonal growth and branching (Fukata *et al.* 2002). CRMP2 also binds to the kinesin-1 light chain subunit and acts as an adaptor for the transport of tubulin heterodimers as well as the actin regulators Sra-1 and WAVE into axonal growth cones (Kawano *et al.* 2005; Kimura *et al.* 2005). Furthermore, CRMP4 physically associates with *in vitro* F-actin (Rosslenbroich *et al.* 2005). In cultured DRG neurons, CRMP1 colocalizes to the actin cytoskeleton (Higurashi *et al.* 2012), and drives actin elongation in lamellipodia formation in cultured epithelial cells (Yu-Kemp *et al.* 2017). These studies indicate that CRMPs can have both positive and negative effects on neuronal protrusion, and most of the biochemical evidence indicates that CRMPs promote actin assembly and MT function. Our results suggest that UNC-33/CRMP has a negative effect on growth cone protrusion and MT entry into growth cones, consistent with the original finding of CRMPs as anti-protrusive factors (Goshima *et al.* 1995; Takahashi *et al.* 1999). The role of UNC-44/ankyrin might be to properly localize UNC-33/CRMP as previously described (Maniar *et al.* 2012). Loss of dorsal F-actin asymmetry and excess protrusion could be a secondary consequence of excess MT accumulation in the growth cone, or could represent independent roles of UNC-33/CRMP.

We have identified three aspects of VD growth cone morphology affected by *unc-5* mutants: excess protrusion; dorsal F-actin accumulation; and restriction of MT + ends from the growth cone. Neither excess MT + ends nor loss of dorsal F-actin polarity alone were sufficient to drive excess protrusion, as *unc-40; unc-5* and *ced-10; mig-2* double mutants display loss of F-actin polarity and excess MT + ends but not excess growth cone protrusion. Thus, an additional mechanism, possibly involving UNC-40, CED-10, MIG-2, and actin nucleators such as Arp2/3, UNC-115/abLIM, and UNC-34/Ena are required to drive protrusion downstream of F-actin polarity and MT + end entry.

Possibly, a dynamic interaction between MTs and actin, mediated by UNC-33/CRMP, controls MT accumulation in the growth cone during repulsive axon guidance mediated by UNC-6/Netrin. The interactions between actin and microtubules in growth cones *in vitro* is well-documented and complex (Dent *et al.* 2011; Coles and Bradke 2015), including the idea that actin retrograde flow removes MTs from the growth cone periphery due to physical linkage to actin undergoing retrograde flow (Lin and Forscher 1995; Lee and Suter 2008; Schaefer *et al.* 2008; Short *et al.* 2016; Turney *et al.* 2016). An intriguing interpretation of our results, based upon those in cultured neurons, is that the UNC-6/Netrin signaling pathway we have described inhibits protrusion by maintaining MT attachment to actin, possibly via UNC-33/CRMP, and thus restriction of MTs from the growth cone. Growth cone dorsal advance could occur by regulated MT entry and interaction with the dorsal leading edge of the growth cone, possibly by interacting with polarized dorsal F-actin. Furthermore, we show that a pro-protrusive function of UNC-40/DCC and the Rac GTPases might act independently of UNC-5 to drive growth cone protrusion, normally at the dorsal leading edge.

### Conclusions

Our results suggest that UNC-6/Netrin signaling coordinates growth cone F-actin accumulation, EBP-2::GFP accumulation, and protrusion to direct growth away from it. UNC-6/Netrin and UNC-5 have a role in polarizing the growth cone, visualized by dorsal F-actin accumulation, resulting in protrusion restricted to the dorsal leading edge. Furthermore, UNC-6/Netrin and UNC-40 stimulate protrusion at the dorsal leading edge, based on the establishment of polarity via UNC-5. This is similar to results in neurons with axons that grow ventrally toward UNC-6/Netrin (e.g. HSN), wherein UNC-6/Netrin and UNC-5 regulate where UNC-40-mediated protrusion can occur in the neuron, in this case ventrally toward the site of UNC-6/Netrin (Kulkarni *et al.* 2013; Yang *et al.* 2014; Limerick *et al.* 2018). These results in axons that grow toward UNC-6/Netrin, along with our results in growth cones that grow away from UNC-6/Netrin, suggest a model of UNC-6/Netrin function involving growth cone polarization coupled with regulation of growth cone protrusion based on this polarity. Recent studies in the vertebrate spinal cord have shown that expression of Netrin-1 in the floorplate is dispensable for commissural axon ventral guidance, (Dominici *et al.* 2017; Varadarajan and Butler 2017; Varadarajan *et al.* 2017; Yamauchi *et al.* 2017) and that contact-mediated interactions with ventricular cells expressing Netrin-1 are more important, consistent with a possible contact-mediated polarity role of Netrin. In any case, several outstanding questions about the polarization/protrusion model presented here remain. For example, how does UNC-6/Netrin result in polarized protrusive activities in the growth cone? Asymmetric localization of UNC-40 and/or UNC-5 is an attractive idea, but UNC-40::GFP shows uniform association of the growth cone margin in VD growth cones and no asymmetric distribution (Norris *et al.* 2014). Also, once established, how is polarized protrusive activity maintained as the growth cone extends dorsally away from the UNC-6/Netrin source? Answers to these questions will be the subject of future study.

---

**Figure 10.**
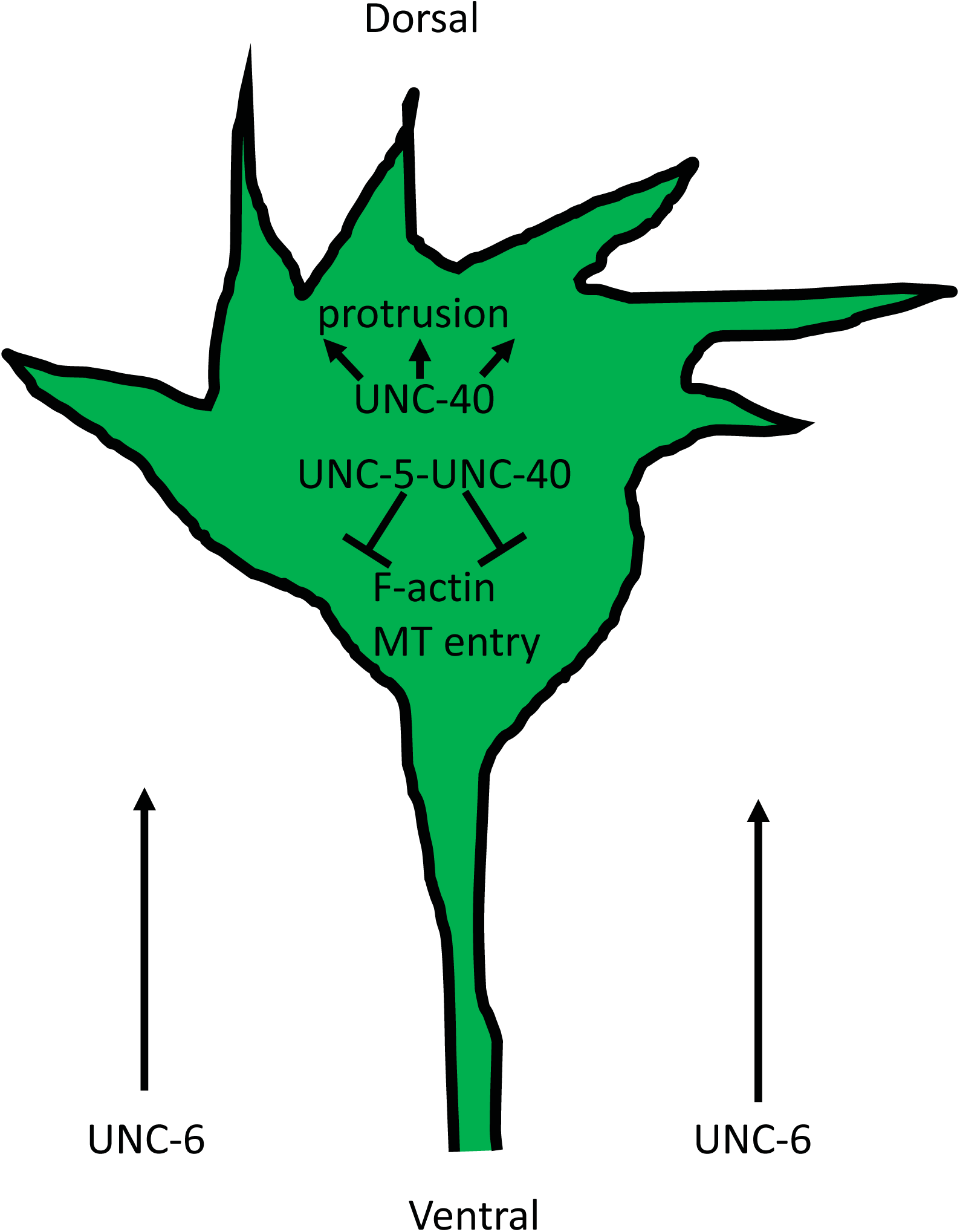
A model of the roles of UNC-5 and UNC-40 in growth cone outgrowth. Our results indicate that UNC-6/Netrin controls multiple, complex aspects of growth cone behavior and morphology during growth away from it. UNC-6 polarizes the growth cone via UNC-5, including F-actin accumulation and protrusion localized to the dorsal leading edge away from the UNC-6 source. UNC-6 also regulates the extent of growth cone protrusion. It inhibits protrusion via UNC-5, possibly by restricting MT + end accumulation in the growth cone. Protrusion can be inhibited independently of MT + ends, possibly via an actin-based mechanism involving the flavin monooxygenases (FMOs). UNC-6/Netrin can also drive growth cone protrusion via UNC-40/DCC. These anti- and pro-protrusive activities of UNC-6/Netrin might act asymmetrically in the growth cone, possible established by the earlier role of UNC-6/Netrin in polarizing the growth cone.

